# Multimodal profiling reveals the centromedian nucleus of thalamus as a dynamic hub orchestrating staged consciousness recovery

**DOI:** 10.64898/2026.05.04.722649

**Authors:** Yulin Zhang, Xiaoyu Wang, Yang Xu, Yuanyuan Yu, Yawen Tan, Hao Wu, Sheng He, Liping Wang, Feng Wang

## Abstract

Consciousness recovery from general anesthesia represents a fundamental brain state transition. The absence of objective, continuous metrics to monitor the shift from unconsciousness to full awareness has left its dynamic, multidimensional nature unresolved, fracturing consciousness theoretical framework and posing clinical risks due to unreliable assessment of emergence. To address this, we developed a multiscale framework integrating AI-driven behavioral analysis, continuous neuro-physiology recording, and whole-brain functional imaging. This decodes recovery into three hierarchical stages, reflex restitution, level restoration, and content re-establishment, each with unique multimodal signatures. We identified heart rate stabilization as a potential noninvasive biomarker for the restoration of consciousness level. Crucially, the centromedian nucleus of thalamus acts as a dynamic hub, actively orchestrating staged whole-brain reconfiguration via stage-dependent network routing. These findings resolve the temporal orchestration of consciousness recovery, ground disparate theories within a unified mechanistic sequence, and open transformative paths for real-time monitoring and neuromodulation in anesthesiology.

**Highlights:**

A multimodal framework decodes consciousness recovery into three hierarchical stages
Recovery stages span from reflex restitution, to level restoration, and finally to content re-establishment
Heart rate stabilization serves as a noninvasive biomarker for consciousness level restoration
The CM paces this staged reconfiguration as a dynamic network hub

**In Brief:** Zhang et al. develop a multiscale framework that delineates consciousness recovery from anesthesia as a three-stage process, identifies heart rate stabilization as a biomarker for the restoration of consciousness level, and reveals the CM as a dynamic hub orchestrating this progression. These findings clarify the temporal orchestration of consciousness recovery, pave the way for real-time monitoring and neuromodulation in anesthesiology and other fields.

## INTRODUCTION

Understanding the neural basis of consciousness remains a paramount scientific challenge.^1–3^ Leading theories, from the Global Neuronal Workspace (GNWT)^4^ to Integrated Information Theory (IIT),^5^ converge on the necessity of large-scale, integrated brain activity for conscious experience.^6^ However, these theories primarily describe correlates of a stable conscious state, lacking a dynamic, mechanistic account of how the brain transitions into and out of that state. A critical unresolved question persists, how does the brain dynamically orchestrate its own reintegration from a fragmented, unconscious state to a coherent, conscious one? The precise temporal processes, governing neural hubs, and continuous biomarkers of this transition remain largely unknown.

This fundamental theoretical gap has direct and serious implications for clinical practice. General anesthetics provides a powerful, reversible model to investigate conscious state transitions.^7,8^ Nevertheless, a stark disconnect exists between this scientific paradigm and clinical reality.^9^ The assessment of consciousness recovery from anesthesia still relies on binary, subjective behavioral indicators such as eye-opening^10–12^ or responses to commands.^11,13–16^ These measures cannot capture the staged, multidimensional nature of neural reactivation across behavioral, autonomic, and network levels. This reliance creates a dangerous diagnostic blind spot in postoperative care, where complications like delayed emergence or covert consciousness may go undetected, potentially compromising patient safety.^17–19^

We therefore hypothesized that consciousness recovery is not a unitary switch but a hierarchically staged process of whole-brain reconfiguration, orchestrated by a specific neural hub. To address this, we developed a unified multiscale framework that integrates AI-driven behavioral analysis, continuous neuro-physiology recording, and whole-brain functional imaging. This approach moves beyond descriptive recovery timeline to establish a mechanistic map linking specific behavioral landmarks to underlying neuro-physiological and network events.

Our framework decodes consciousness recovery into three hierarchical stages, reflex restitution, level restoration, and content re-establishment, each defined by unique multimodal signatures. We identified heart rate stabilization as a noninvasive biomarker for the critical restoration of consciousness level and, crucially, pinpointed the centromedian thalamic nucleus (CM) as a dynamic hub that actively orchestrates this staged whole-brain reconfiguration through stage-dependent network routing. These findings transform consciousness recovery from a phenomenological observation into a decodable process. The identified hub and biomarker offer a fundamental principle and toolset for illuminating state transitions in anesthesia, sedation, and disorders of consciousness, bridging consciousness theory with systems neuroscience, anesthesiology and neurology, and opening avenues for real-time monitoring and targets neuromodulation.

## RESULTS

### Integrated behavioral-cardiovascular signatures characterize staged consciousness recovery

To precisely track the dynamics of consciousness recovery after general anesthesia, we first developed an integrated platform for synchronous collection and multimodal analysis of behavioral and cardiovascular data (Figures S1A and S1B). Using a computational pipeline, we extracted 3D kinematic features and movement sequences from videos by our hierarchical behavioral analysis framework (HBAF)^20^ and processed cardiovascular signals into continuous blood pressure and heart rate traces. These aligned data streams generated a unified, synchronized dataset for multimodal analysis (Figure S1C). This approach identified 17 movements categorized into 6 behavioral clusters, including locomotion, exploration, maintenance, nap, anesthetic posture, and post-anesthetic ataxia (Figures S1D–S1F; Table S1). Further, we investigated temporal recovery patterns during the post-anesthesia (PA) period. Continuous 60-minute characteristics of movement sequence, as well as key kinematic metrics, such as nose height, nose speed, body curling, body length, back height, and distance (Figures S2A–S2C), revealed a time-dependent behavioral resurgence in PA mice that progressively approximated wakefulness (WF) patterns (Figure 1A). Behavioral repertoire analysis conducted in 10-minute intervals showed that PA mice initially exhibited profoundly different behavioral patterns with WF mice, characterized by reduced locomotion and exploration alongside increased post-anesthetic posture and ataxia. Notably, these disruptions persisted after recovery of righting reflex (RORR), gradually normalized to WF-like states approximately 40 minutes post-isoflurane discontinuation (Figures 1B, 1C, S2D, and S2E).

**Figure 1.**
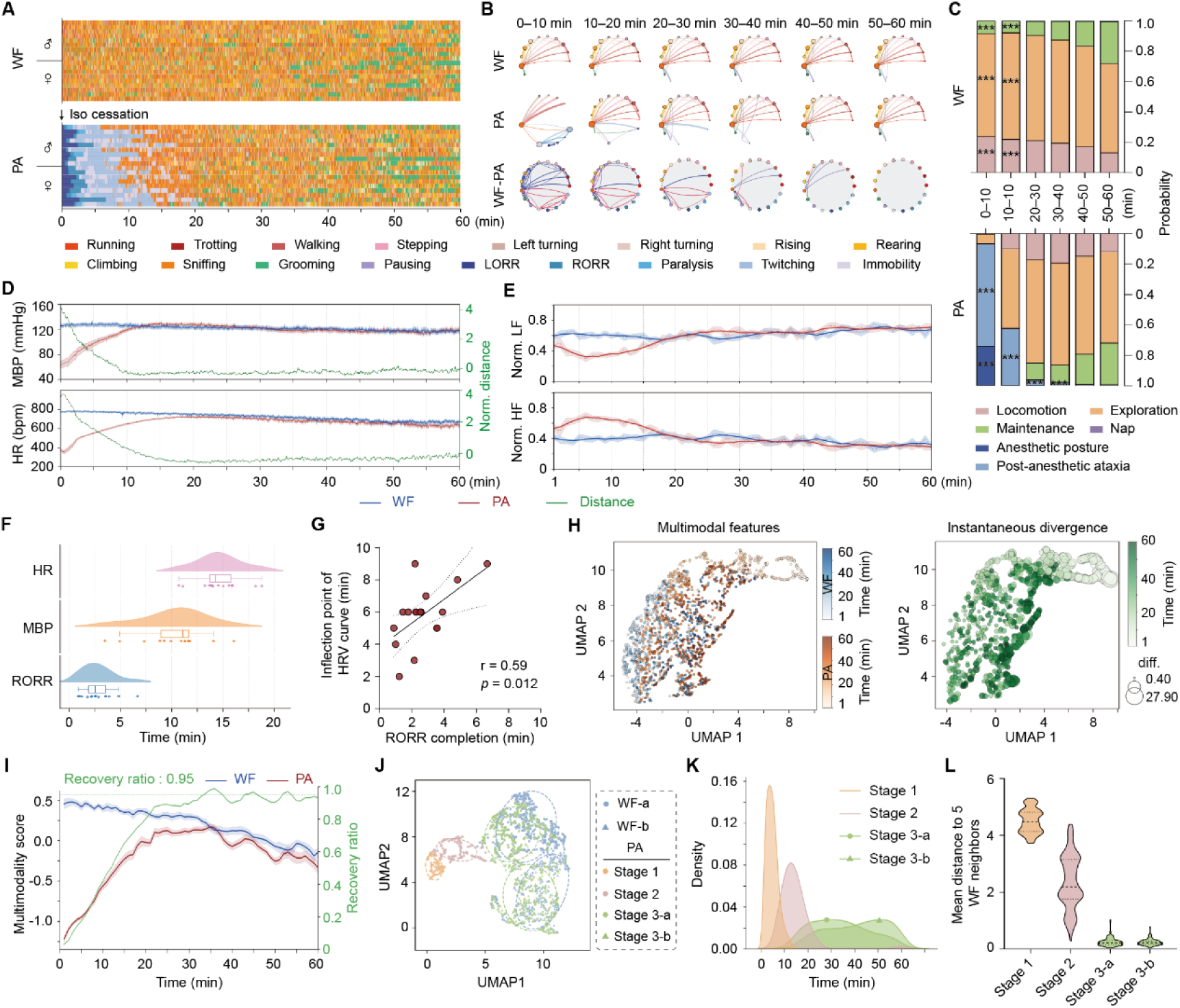
Multimodal tracking of graded consciousness recovery after anesthesia (A) Continuous ethograms during wakefulness (WF, top) and post-anesthesia (PA, bottom) periods (n = 9 male and 9 female mice). (B) Movement transition across 10-minute epochs in WF (top) and PA (middle), and the difference between WF and PA matrices (bottom). (C) Temporal probability distributions of behavioral clusters in WF and PA groups. Statistical analyses are performed using multiple Mann-Whitney tests with Bonferroni–Dunn correction for multiple comparisons. (D) Dynamics of mean blood pressure (MBP, top) and heart rate (HR, bottom) during WF and PA periods (left Y-axis), and Dynamic Time Warping (DTW) distance between WF and PA sequences (right Y-axis). (E) Dynamics of normalized low frequency (top) and high frequency (bottom) HRV power. (F) Recovery latencies for the RORR, MBP, and HR. (G) Correlation between RORR and HRV inflection timing. (H) UMAP projection of multimodal features for WF (blue) and PA (orange) groups (left). Instantaneous divergence of PA states from WF baseline over time (right). (I) Multimodality score evolution (left Y-axis) and recovery ratio (right Y-axis). (J) Spectral clustering of PA stages within UMAP space, with 95% confidence ellipses. The WF group is shown for reference. (K) Temporal density distribution of PA stages. (L) Mean distance from each PA stage to its nearest neighbors in the WF baseline. Data are presented as shows mean (A), mean ± s.e.m. (I), median with quartiles (F and L), and probability density (K). \*\*\**P* < 0.001, \*\**P* < 0.01, \**P* < 0.05. Statistical details are presented in Table S2.

Cardiovascular parameters followed a similar recovery trajectory. Blood pressure (BP), including diastolic (DiaBP), systolic (SysBP), and mean (MBP), as well as NN interval and heart rate (HR), displayed a gradual normalization toward WF levels (Figures 1D, S3A, and S3B). Dynamic Time Warping (DTW) distance quantification revealed an increasing similarity of cardiovascular parameters between PA and WF states over time (Figure 1D). Full recovery of BP and HR lagged behind RORR by approximately 10 and 15 minutes, respectively (Figure 1F). Heart rate variability (HRV) parameters exhibited dynamic biphasic recovery patterns, with low-frequency (LF) power initially decreasing before gradual increasing, while high-frequency (HF) power demonstrated an initial rise followed by a decline, ultimately stabilizing at WF-comparable levels (Figure 1E). This pattern suggested an initial reactivation of the parasympathetic system, followed by a rebalancing between sympathetic and parasympathetic activities. The inflection points in HRV closely correlated with the completion time of RORR (Figure 1G), indicating that RORR represents a pivotal neurophysiological transition where central nervous system reactivation enables both reflex recovery and autonomic recalibration.

Uniform Manifold Approximation and Projection (UMAP) of high-dimensional behavioral-cardiovascular data revealed a clear separation between WF and early PA stages, with PA trajectories progressively converging toward WF intervals (Figure 1H, left). The instantaneous divergence between PA and WF decreased over time, reflecting a continuous recovery process (Figure 1H, right). We developed a multimodal score that integrated behavioral and cardiovascular metrics, which rapidly increased during the first 20 minutes of the PA phase before gradual convergence with WF trends.

Recovery ratio analysis indicated a 95% return to WF baseline at approximately 35 minutes into the PA phase (Figure 1I). Spectral clustering identified three discrete recovery stages, each with unique behavioral-cardiovascular profile and temporal distributions (Figures 1J and 1K), showing a progressive consciousness convergence (Figure 1L). Stage 1 featured basic reflex recovery accompanied by autonomic instability and significant ataxia (Figures 1A–1D and S2A–S2D). Stage 2 marked the initiation of pronounced cardiovascular recovery (Figures 1D–1F) and environmental exploration characterized by behavioral disinhibition (Figures 1A, 1C, S2D, and S2E). Stage 3 demonstrated behavioral organization (Figures 1A–1C, S2D, and S2E) and cardiovascular dynamics (Figures 1D, 1E, S3A, and S3B) that approximated WF levels, indicating near-complete functional reintegration. This stage was further subdivided based on alignment with early (higher valence) and later (lower valence) WF baseline segments (Figure 1J), reflecting natural fluctuations in conscious states (Figure 1I). Recovery trajectories remained consistent across sexes and individuals (Figures S3C and S3D). Electroencephalography (EEG) analysis revealed a shift from slow-wave dominance toward a balanced, wake-like power spectrum (Figures S3E–S3G), temporally coupled with behavioral recovery, underscoring the integrated nature of consciousness restoration. These findings established that consciousness recovery from general anesthesia is a staged process characterized by the coordinated, time-dependent recurrence of behavioral-cardiovascular features, thereby providing a quantitative framework to investigate its underlying neural mechanisms.

### Defensive responses to visual threats indicate recovery of consciousness level

Consciousness encompasses two key dimensions: arousal (level of consciousness) and awareness (content of consciousness).^21,22^ Arousal necessitates integrated sensorimotor coordination and appropriate emotional responses.^23,24^ Having established a staged recovery framework, we next sought to identify stage-specific behavioral landmarks that could reliably track the restoration of consciousness level. Given that looming stimuli evoked innate defensive responses that engage these capacities, we hypothesized that defensive responses to visual threats (DRVT) could effectively indicate the recovery of consciousness levels. We established a paradigm to elicit and quantify DRVT (Figure 2A). Ethograms analysis revealed that the probability of defensive responses was exquisitely dependent on the level of consciousness, gradually recovering from complete absence immediately after the RORR (PR0) and at 5 minutes post-RORR (PR5), to partial response at 10 minutes post-RORR (PR10), and approaching WF levels by 15 minutes post-RORR (PR15) for both sexes (Figures 2B–2D and S4A). The quality of these responses, assessed through flight response latency, return speed and time to refuge, was similarly compromised during early recovery (PR10) but largely restored to WF-level performance by PR15 (Figures 2E, 2F, and S2B). This staged recovery extended beyond defensive behaviors to a restructuring of the behavioral repertoire, as movements probability and distribution of behavioral clusters progressively aligned with WF states over time (Figures 2G, 2H, and S4C).

**Figure 2.**
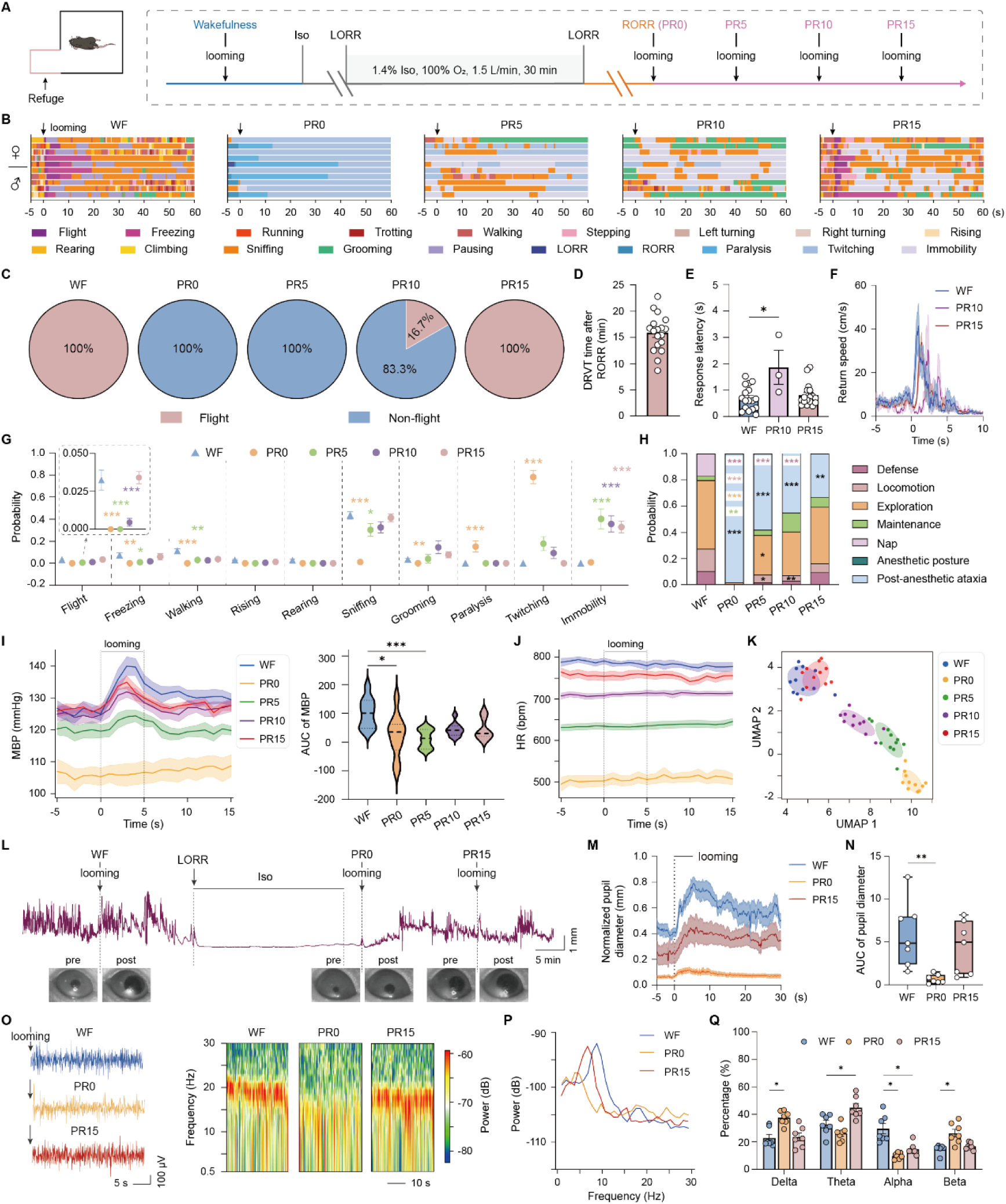
Recovery of defensive responses marks consciousness level progression. (A) Experimental schematic of true refuges (1T) behavioral apparatus and testing paradigm. (B) Representative ethograms following looming stimuli presentation across consciousness timepoints (WF, PR0, PR5, PR10, and PR15). Arrowheads indicate looming stimuli delivery. (C) Flight probabilities across recovery timepoints (n = 7 male and 11 female mice). (D) DRVT recovery latency after RORR. (E and F) Response latency (E) and return speed (F) among WF, PR10, and PR15. (G and H) Probabilities of movements (G) and behavioral clusters (H) across timepoints. (I) Mean arterial pressure (MBP) dynamics (left) and integrated response magnitudes (right) following looming (n = 7 male and 11 female mice). (J) Heart rate (HR) trajectories following looming stimuli. (K) UMAP projection of multimodal features across recovery timepoints (n = 5 male and 6 female mice). (L) Representative pupil diameter dynamics after looming stimuli during WF, anesthesia, and PA periods. (M) Normalized pupil responses to looming stimuli (n = 4 male and 3 female mice). (N) Integrated pupillary response magnitudes. (O) Representative EEG traces (left) and spectrograms after looming stimuli. (P) Mean EEG power spectral density profiles. (Q) Quantitative EEG band power comparisons (n = 4 male and 3 female mice). Data are presented as percentage distributions (C), mean (H and P), mean ± s.e.m. (D–G; I, left; J, M, N, and Q), and median with quartiles (I, right). Statistical analyses are performed using Kruskal-Wallis test with Dunn’s multiple comparisons test (E and I), one-way ANOVA with Dunnett’s post hoc test (N), and Mann-Whitney tests with Bonferroni–Dunn correction for multiple comparisons (G, H, and Q). \*\*\**P* < 0.001, \*\**P* < 0.01, \**P* < 0.05. Statistical details are presented in Table S3.

Synchronous recordings of cardiovascular parameters revealed clear recovery patterns. The typical pressor response to looming threat was diminished or absent at PR0 but gradually returned to the pronounced response seen in the WF states (Figures 2I and S4C–S4E). Although tachycardic responses were not observed, baseline HR values displayed a similar staged recovery, reaching WF levels by PR15 (Figures 2J, S4C, and S4F). When projecting these integrated behavioral-cardiovascular data into UMAP space, we captured the entire recovery trajectory, showing a distinct separation among consciousness levels and a time-dependent convergence of PR states toward WF (Figure 2K). Critically, integrated visual threat features were fully regained by PR15 (Figure 2K), establishing the coordinated recovery of DRVT as a functional landmark for consciousness level restoration.

To systematically assess functional recovery beyond behavioral-cardiovascular levels, we employed complementary physiological assays. While basic motor tone sufficient for the righting reflex was present at PR0 (Figures S4G–S4J), coordinated motor function and nociceptive processing remained impaired until PR15 (Figures S4K–S4M). This dissociation indicates that spinal reflex return precedes complex sensorimotor integration. Autonomic and cortical functions showed parallel, graded recovery. Pupillometry, a readout of autonomic arousal,^25^ showed nearly absent looming-evoked dilation at PR0 (Figure 2L), but significant recovery by PR15 (Figures 2L–2N), while EEG transitioning from delta-dominated slow-wave activity at PR0 to a balanced, wake-like spectral pattern with increased higher-frequency power by PR15 (Figures 2O–2Q). Furthermore, this staged multimodal recovery sequence is invariant to anesthetic mechanism, as demonstrated by identical DRVT restoration dynamics following propofol-induced anesthesia (Figures S5A–S5E). Thus, DRVT recovery marks the critical transition to a conscious state characterized by reintegrated autonomic control, cortical processing, and purposeful motor execution.

Together, these results establish DRVT as a reliable behavioral landmark for the restoration of consciousness level and identify HR stabilization as a tightly coupled physiological signature of this transition. This prompted us to evaluate the translational potential of HR stabilization as a noninvasive biomarker for tracking the recovery of consciousness level. Analysis of postoperative HR records from a large clinical cohort^26^ revealed that abnormal HR recovery (defined as a deviation of >10% from the preoperative baseline) at the defined emergence timepoint was significantly associated with an increased incidence of postoperative adverse events (Figures S6A–S6I). Notably, cardiovascular complications exhibited the strongest association with abnormal HR recovery patterns (Figure S6G). This translational evidence reinforces the potential of HR stabilization to serve as an objective and physiologically grounded biomarker for consciousness recovery in clinical.

### Complex defensive responses signify recovery of consciousness content

To investigate the recovery of consciousness content following general anesthesia, we designed a cognitively demanding multi-refuge defensive response (MRDR) paradigm that requires higher-order cognitive processing. Mice were subjected to two behavioral configurations: a simple ipsilateral true-fake refuge (ips T-F) setup, which tested basic refuge discrimination, and a complex 1-true-3-fake refuge (1T-3F) setup, which assessed spatial awareness and decision-making under threat (Figure 3A). The percentage of mice exhibiting defensive responses to looming stimuli recovered over time, but the recovery trajectories were significantly affected by cognitive demands. In the simple ips T-F condition, response rates approached WF levels by PR15 (Figure 3A). However, in the more complex 1T-3F condition, recovery was slower, with responses remaining suboptimal even at 20 minutes post-RORR (PR20) (Figure 3A). Spatial analysis of post-threat positions and trajectories in the 1T-3F condition revealed disorganized movement patterns at PR10 and PR15. By PR20, however, trajectories became direct and optimized, resembling WF patterns (Figure 3F), indicating that the recovery of consciousness content is a staged process that is sensitive to task complexity (Figure 3A). Moreover, flight initiation latency following RORR was significantly longer in the cognitively demanding 1T-3F condition compared to the simpler ips T-F condition (Figure 3B). Initial escape strategies also varied: in the simple condition, 85% of mice used direct flight strategies, while in the complex condition, only about 60% selected the correct refuge on their first attempt (Figure 3C). The time to successfully flight into the refuge was delayed in the complex condition (Figure 3D), and escape accuracy improved gradually, reaching 100% in the ips T-F condition by PR15, but remaining suboptimal in the 1T-3F condition even at PR20 (Figure 3E), These recovered patterns were consistent across both sexes in both conditions (Figures S7A–S7D).

**Figure 3.**
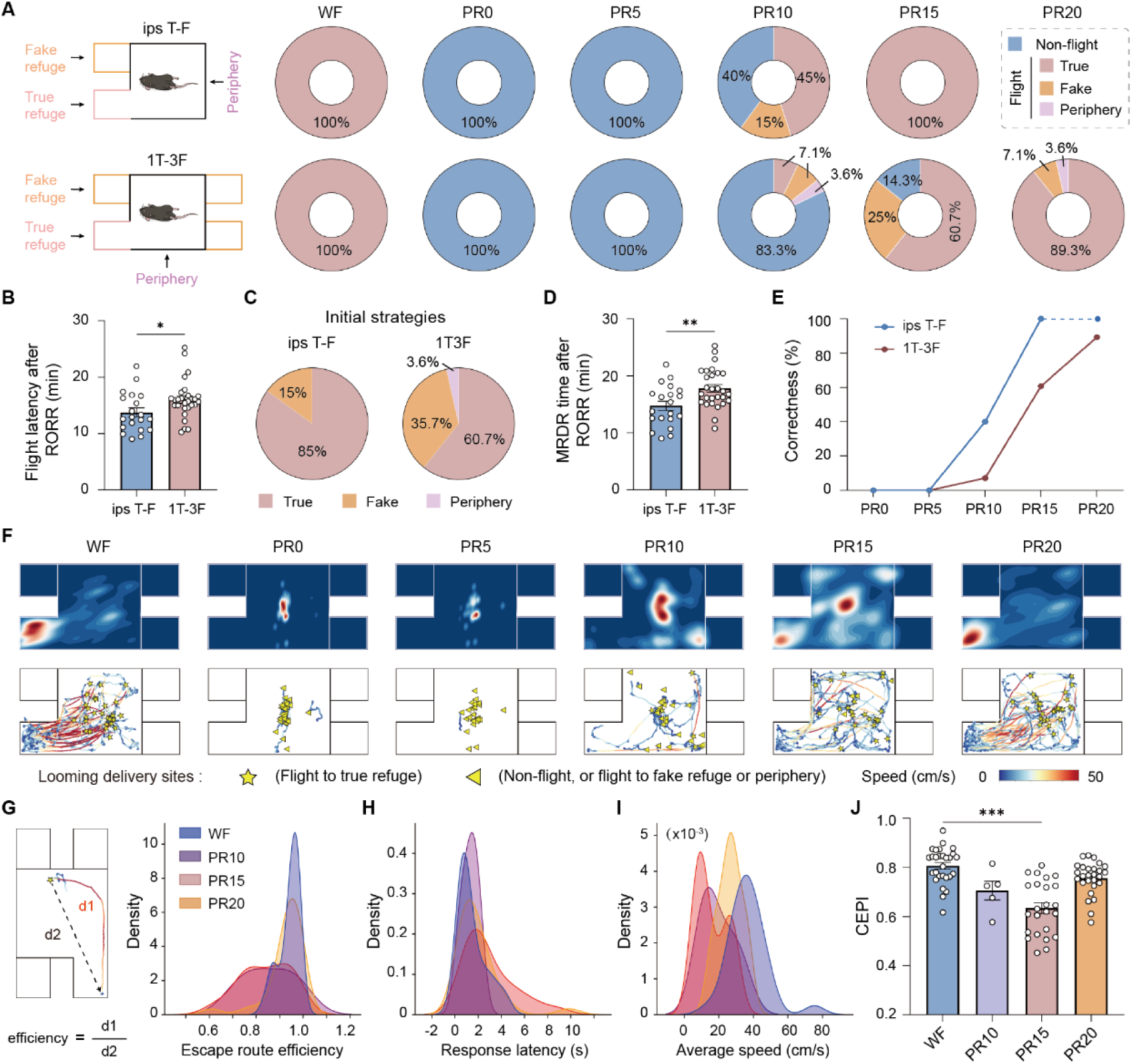
Complex defensive responses reflect consciousness content restoration. (A) Apparatus schematics for ipsilateral true-fake refuges (ips T-F, top) and 1-true-3-fake refuges (1T-3F, bottom) behavioral apparatus, with flight probabilities across recovery timepoints (n = 20 mice (10 per sex) in ips T-F, n = 29 mice (15 male and 14 female) in 1T-3F). (B) Flight initiation latencies post-RORR across behavioral configurations. (C) Initial escape strategy distributions. (D) MRDR latencies after RORR. (E) Population escape accuracy over time. (F) Spatial density distributions (top) and individual trajectories (bottom) following looming stimuli in 1T-3F configuration. (G) Escape route efficiency calculation (left) and probability distributions across time points (right). (H and I) Response latency (H) and return speed (I) distributions across PR time. (J) Composite escape performance index (CEPI) comparisons. Data are presented as percentage distributions (A, C, and E), mean ± s.e.m. (B, D, and J), and probability density (G–I). Statistical analyses are performed using unpaired *t*-test (B and D) and Kruskal-Wallis test with Dunn’s multiple comparisons test (J). \*\*\**P* < 0.001, \*\**P* < 0.01, \**P* < 0.05. Statistical details are presented in Table S4.

Escape route efficiency, which combines path directness and success, increased progressively across PR time points, achieving WF-like levels by PR20 (Figures 3G and S7E). Defensive kinematics, including response latency, return speed, and time in refuge, largely return to math WF profiles at this time (Figures 3H, 3I, and S7F–S7H), indicating that decision-making and motor execution accelerated as consciousness content recovered by PR20. We computed a composite escape performance index (CEPI) that demonstrated achievement of WF-level performance by PR20 (Figure 3J). Behavioral sequences and the movement and clusters probabilities gradually aligned with the WF distribution (Figures S7I–S7K). In summary, these findings indicate that MRDR provides a sensitive, continuous measure of consciousness content recovery, effectively capturing the restoration of awareness during the staged recovery process following anesthesia.

### Functional networks undergo spatiotemporal reorganization during consciousness recovery

To explore the whole-brain spatiotemporal dynamics underlying staged consciousness recovery from anesthesia, we utilized an awake functional magnetic resonance imaging (fMRI) paradigm in head-fixed mice (Figure 4A) and identified three distinct whole-brain states through dynamic functional connectivity (FC) analysis (Figure S8A). State 1 exhibited low global connectivity with fragmented sensory-thalamic-limbic architecture; State 2 displayed moderate connectivity with integrated association cortices and thalamo-hippocampal circuits; State 3 was characterized by high global connectivity and widespread yet simplified co-activation across sensorimotor and subcortical neuromodulatory regions (Figures S8A, S8D, and S8E). During consciousness recovery, these states evolved nonlinearly (Figures S8B and S8C), initiated with a brief, unstable period of high integration and hub dominance, progressing through a phrase of meta-stability with fluctuating transition entropy, and finally stabilizing into a flexibly reconfigured network (Figure 4B). The finding that identical FC states exhibited time-varying transition probabilities (Figure 4C) further indicates a dynamic temporal reorganization underlying this staged recovery.

**Figure 4.**
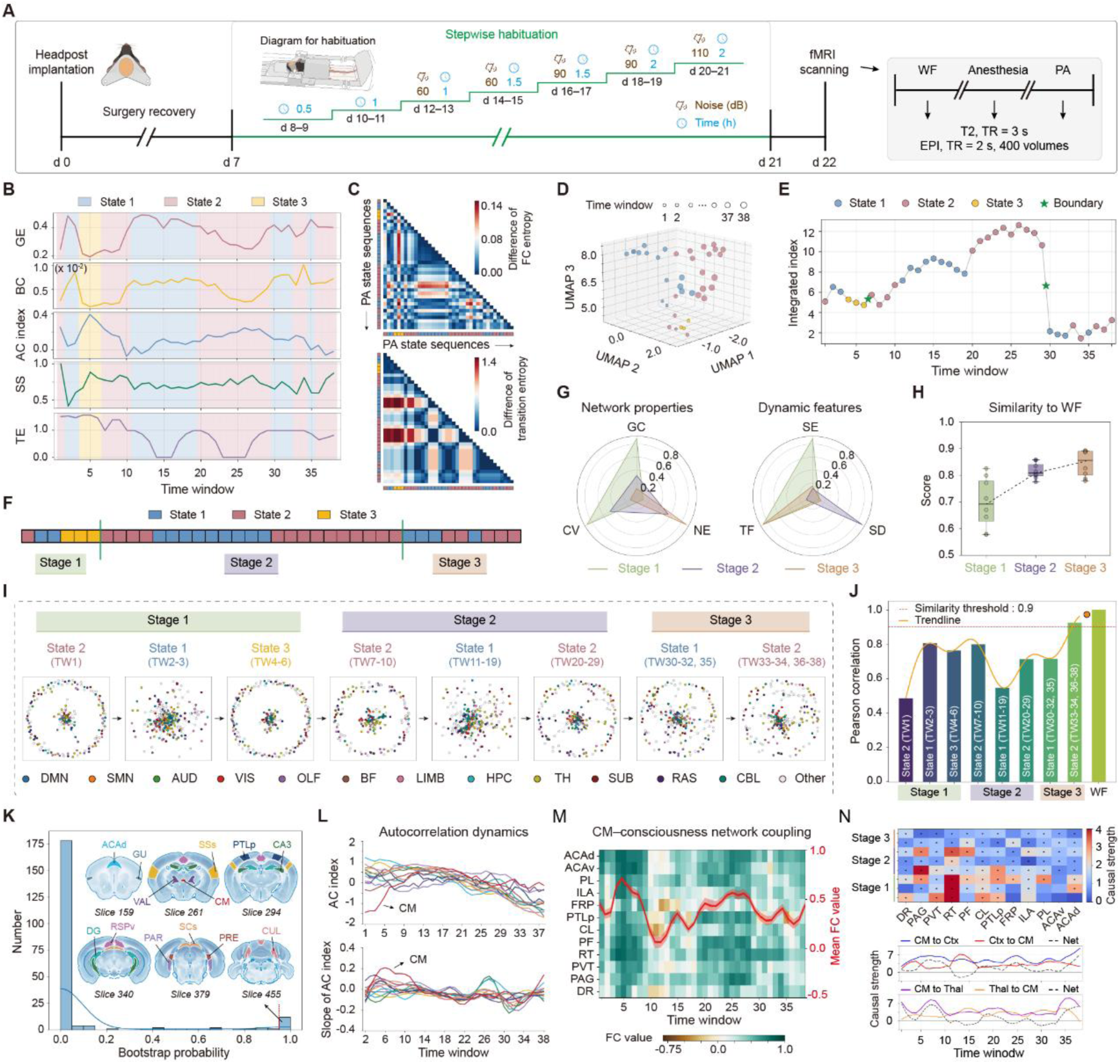
Spatiotemporal reorganization of functional networks during consciousness recovery. (A) Experimental timeline with headpost design and habituation protocol, as well as fMRI scanning procedure. (B) Dynamic trajectories of nodal and global network features (GE, global efficiency; BC, betweenness centrality; AC, autocorrelation; SS, state stability; TE, transition entropy). (C) Difference of functional connectivity (FC) entropy (top) and state transition entropy (bottom). (D) UMAP projection of integrated features across timepoints. (E) Hierarchical recovery sequence of functional networks. (F) Stages delineation through change-point analysis. (G) Radar charts of network properties (left; GC, global connectivity; CV, connectivity variability; NE, network efficiency) and dynamic features (right; SE, state entropy; TF, transition frequency; SD, state duration) across stages. (H) FC similarity progression to WF baseline. (I) Evolution of functional connectome across stages. (J) Pearson correlation of FC matrix between each stage and WF. Orange curve and dot represent trend in similarity and predict subsequent values across stages, respectively. Red line indicates a designed recovery threshold (0.9). (K) Hub region stability analysis based on bootstrap resampling (outset), and the spatial distribution of high-stability hubs (inset). The blue curve represents kernel density estimation. (L) Temporal profiles of the autocorrelation index (top) and its instantaneous slope (bottom) for selected high-stability hub regions. (M) Dynamic FC patterns between the centromedian nucleus of thalamus (CM) and other consciousness-related regions across recovery stages. (N) Dynamic heatmap of significant Granger causal influences from the CM to other regions across stage (top; asterisks denote p<0.05). Temporal evolution of the net directional causal flow (CM→target minus target→CM) between the CM and the cortex (Ctx) or thalamus (Thal) aggregate regions (bottom). Black dashed lines highlight the net driving effect. Data were from 4 male and 4 female mice. Data are presented as mean (B, G, L, and N), mean ± s.e.m. (M), and median with quartiles (H). Anatomic abbreviations are shown in Table S5.

Projecting the multi-dimensional features of these three states, including information-theoretic metrics, network connectivity, and state transitions, into UMAP space revealed three distinct clusters (Figure 4D). These neural clusters appear to correspond to the three behavioral-cardiovascular stages of recovery defined earlier (Figures 4E and 4F), suggesting the staged recovery observed in behavior and physiology is intrinsically mirrored by, and likely driven by, a parallel reorganization of global brain network dynamics. Each recovery stage was not a monolithic brain state but a stage-specific sequence or combination of these fundamental network states (Figures 4F and 4G). This spatiotemporal progression was marked by a significant increase in whole-brain functional connectivity similarity to the awake baseline, exceeding 85% in Stage 3 (Figure 4H). Blood-oxygen-level-dependent (BOLD) fluctuations and activation patterns provided further validation (Figures S9A–S9C). Furthermore, the network reconfiguration trajectory—from initial fragmentation to transient resegregation and ultimately to optimized integration (Figures 4I and 4J), closely paralleled autonomic fluctuations in heart rate variability (Figure 1E), suggesting a deep coupling between the brain’s network restructuring and systemic autonomic rebalancing during consciousness recovery.

To identify neural drivers of this reorganization, we mapped the whole-brain recovery initiation sequence through autocorrelation changes and identified 69 regions with the earliest recovery onset, predominantly in cortical and thalamic areas (Figures S9D and S9E). High-stability hubs (scores > 0.95) included isocortical (e.g., supplemental somatosensory area and anterior cingulate area), thalamic (e.g., ventral anterior-lateral complex and centromedian nucleus), and midbrain (e.g., superior colliculus) regions (Figure 4K). Analysis of the dynamic autocorrelation index revealed that the centromedian nucleus of thalamus (CM) activity had both a high initial value and steep recovery slope during Stage 1, indicating a uniquely heightened and rapidly evolving state compared to other regions (Figure 4L). Dynamic FC between CM and other consciousness-related nuclei^6,27–32^ showed a characteristic stagewise progression, widespread and high-amplitude coupling in Stage 1, selective disengagement and reconfiguration in Stage 2, and an optimized, core-like connectivity profile in Stage 3 (Figure 4M). Furthermore, analysis of Granger causal influence^33^ from CM to these regions across stages demonstrated that CM consistently served as a dominant driver throughout consciousness recovery process. This driving effect exhibited a distinct temporal pattern: strong early effects on thalamus and midbrain nuclei (Figure 4N, top), followed by progressively strengthened influence on prefrontal cortical nuclei in later stages (Figure 4N, bottom). Given its prominent dynamic activity profile and stage-dependent network connectivity, CM emerged as a leading candidate core hub governing network dynamic, prompting further investigation into its causal role in orchestrating staged recovery.

### The centromedian nucleus of thalamus orchestrates staged consciousness recovery

Building on the identification of CM as a stable network hub, we next employed causal interventions to determine whether it actively paces progression through the defined recovery stages. Calcium activity of CM showed an oscillatory decline following the loss of the righting reflex under anesthesia (Figures 5A and 5B). During the PA period, CM activity began to recover prior to the initiation of RORR and increased rapidly before RORR completion (Figures 5C and 5D). By PR0, CM activity had recovered to a level comparable to WF and continued to gradually increase until matching the full WF state (Figures 5E and 5F). Chemogenetic activation of the CM immediately after anesthesia accelerated all stages of consciousness recovery (Figures 5H–5J). It reduced RORR latency (Figures 5H and S10A), shortened the recovery of defensive behaviors in both simple and complex environments (Figures 5H and S10B–S10E), decreased the time to reach maximum behavior entropy (Figures 5I and 5J), and speeded up the reorganization of spontaneous behavior (Figures S10F and S10G). Conversely, chemogenetic inhibition of the CM delayed recovery stages (Figures 5K–5M). It prolonged RORR (Figures 5K and S10H), delayed recovery of DRVT and MRDR (Figures 5K and S10I–S10L), and slowed down spontaneous behavioral reorganization (Figures 5L, 5M, S10M, and S10N). These regulation effects are consistent with previous evidence that CM activity correlates with conscious perception in human.^34^

**Figure 5.**
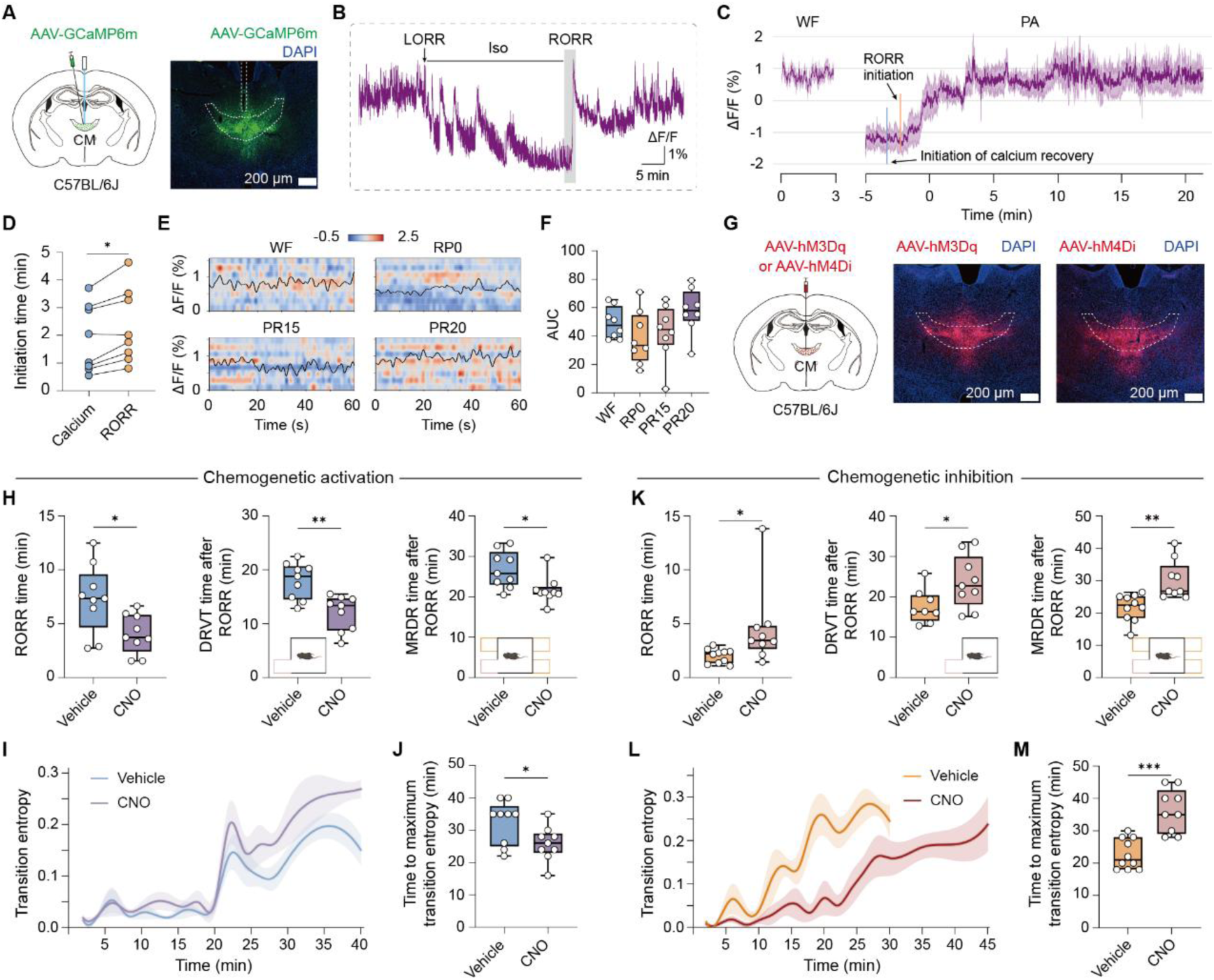
Centromedian nucleus of thalamus orchestrates staged consciousness recovery. (A) Viral injection strategy and representative GCaMP6m expression in the CM. (B) Representative calcium transients throughout the WF-anesthesia-PA period. (C) Group-level calcium signals during WF and PA periods (n = 4 male and 4 female mice). Vertical lines indicate the mean time of CM calcium recovery onset (blue) and RORR onset (orange). (D) Quantification of initiation time for calcium recovery and RORR. (E and F) Heatmap (E) and AUC (F) of CM calcium in 1-minute epoch across stages. (G) Chemogenetic targeting strategy and expression verification in the CM. (H to J) Recovery parameters following CM activation (n = 9 mice (5 male and 4 female) in per group for both 1T and 1T-3F). RORR latency (H, left), defensive behavior recovery in simple (H, middle) and complex (H, right) conditions, behavioral entropy dynamics (I), and peak entropy timing (J). (K to M), Recovery parameters following CM inhibition (For 1T, n = 8 mice (4 per sex) in vehicle and 9 mice (5 male and 4 female) in CNO; for 1T-3F, n = 10 mice (5 per sex) in vehicle and 9 mice (5 male and 4 female) in CNO). RORR latency (K, left), defensive behavior recovery in simple (K, middle) and complex (K, right) conditions, behavioral entropy dynamics (L), and peak entropy timing (M). Data are presented as mean ± s.e.m. (C, F, I, and L) and median with quartiles (F, H, J, K, and M). Statistical analyses are performed using paired *t*-test (D), unpaired *t*-test (H, J, K, and M), and one-way ANOVA with Dunnett’s post hoc test (F). Scale bars, 200 µm (A and G). \*\*\**P* < 0.001, \*\**P* < 0.01, \**P* < 0.05. Statistical details are presented in Table S6.

To test the specificity of this pacemaking function, we also examined two other candidates. The superior colliculus (SC) is a key subcortical hub involved in sensory integration^35,36^ and was identified as one of the high-stability hubs (Figures 4K and 4L). Calcium imaging indicated a gradual recovery of SC following anesthesia (Figures S11A and S11B), with DRVT observed during WF and at PR15, but absent at PR0 (Figures S11B–S11D). This suggests that SC responsiveness to visual threats require appropriate levels of consciousness. However, optogenetic activation of the SC, either before or after RORR, failed to accelerate consciousness restoration (Figures S11E–S11I), indicating that SC engagement does not actively pace consciousness recovery. We also investigated the perifornical hypothalamic area (Pef), a well-known wake-promoting region.^22,37^ Consistent with the previous studies,^38^ calcium activity in the Pef sharply increased during RORR (Figures S11J–S11M), and its activation accelerated RORR (Figures S11N–S11P). However, activating the Pef did not influence DRVT responses (Figure S11Q), indicating selective facilitation of reflexive function without promoting higher-order consciousness recovery.

Collectively, these findings establish the CM as a crucial hub that dynamically orchestrates staged consciousness recovery across all landmarks, from basic arousal to higher-order awareness, distinguishing it from both sensory integration centers and traditional wake-promoting regions.

## DISCUSSION

Our study moves beyond the binary switch model of consciousness to reveal a CM orchestrated, staged cascade of whole-brain reconfiguration. The integrated framework we used does not merely describe recovery; it provides the first systematic decoding of the process, delivering a multiscale mechanistic map that links specific behavioral landmarks (RORR, DRVT, MRDR) to underlying autonomic, network, and circuit events. We thereby establish a detailed, quantitative landscape of recovery as a structured three-stage process, reflexive arousal, perceptual integration, and cognitive restoration, and identified heart rate stability as a potential robust, noninvasive biomarker for the restoration of consciousness level which accurately tracks the critical transition to perceptual integration. Most importantly, we revealed that the CM acts as a core dynamic hub, actively orchestrating this staged whole-brain reprogramming.

Our findings ground prominent consciousness theories within a dynamic, mechanistic framework. The early perceptual integration stage, indexed by cardiovascular stabilization and recovery of innate defensive response (DRVT) prior to complex cognitive access, aligns with theories emphasizing phenomenal awareness^39–41^ and interoceptive inference.^42–44^ This stage may reflect the initial establishment of a coherent perceptual scene. This is supported by the integration within posterior cortical regions, as emphasized by IIT,^5,6,41,45^ and is also consistent with interoceptive Predictive Processing theories, where stabilization of bodily state inference forms the bedrock of conscious experience.^6,46^ Mechanistically, this stage is characterized by CM-driven, widespread high-amplitude thalamocortical coupling, scaffolding foundational integration. The subsequent cognitive restoration stage, marked by complex conscious perception (MRDR) and executive functions, corresponds to the onset of widespread cognitive access and meta-representational capacities. This stage aligns with GNWT^4,6^ and Higher-Order Theories,^6,47,48^ as the network reorganizes into an optimized, flexible core-periphery architecture supporting global information broadcasting. Here, the network reorganizes into an optimized, flexible core-periphery architecture, and CM’s causal influence shifts toward prefrontal cortical nuclei to support global information broadcasting. Critically, CM is not merely a general arousal nucleus (like Pef) nor a passive sensory integrator. Instead, it functions as a dynamic, stage-dependent routing hub. Early on, it drives subcortical and thalamic circuits to set a global integrative tone, converges with the Dendritic Integration Theory.^49,50^ Later, it optimizes prefrontal connectivity to support higher cognition. Thus, CM acts as a conductor orchestrating the staged “symphony” of whole-brain reprogramming. Our work thereby proposes a unifying perspective: consciousness recovery is a CM-orchestrated, staged process of dynamic whole-brain reconfiguration, integrating aspects highlighted by disparate theories,^6,51–54^ with CM serving as the key dynamic node linking interoceptive stability to global cognitive access.

This paradigm resolves previously puzzling observations, such as the dissociation where basic reflexes return long before purposeful action,^55^ and the lag of cardiovascular stabilization behind motor reflex recovery.^56^ These can now be understood as natural manifestations of distinct physiological stages. Our multi-modal staging toolset, particularly the heart rate stability biomarker, provides the anesthesiology and neurocritical care communities with a crucial objective, continuous metric to track consciousness depth, addressing a decades-old diagnostic void. For researchers, the framework offers a new lens to re-examine data from anesthesia studies, psychedelic states, or brain injury recovery, where similar transitions may occur.

Clinically, our findings enable a paradigm shift from passive observation^11^ to active monitoring and intervention. Continuous HR monitoring emerges as a potential, noninvasive tool for real-time assessment, especially in non-communicative patients. Furthermore, we identify CM as a promising therapeutic target for neuromodulation. Its causal role of CM in pacing recovery trajectories aligns with previous evidence that central thalamic stimulation restores consciousness signatures in a nonhuman primate model.^57^ Neuromodulation techniques like deep brain or transcranial magnetic stimulation targeting CM could potentially accelerate consciousness recovery, particularly in cases of delayed emergence. Crucially, the CM-driven staging mechanism, if found to be universal across anesthetic agents and other states of consciousness depression, would position CM as a fundamental node for broad therapeutic modulation. Furthermore, the framework implies that pathologies of consciousness, such as recovery stagnation in minimally conscious states, may reflect a failure to progress through these defined stages. This directly suggests that CM-targeted neuromodulation might not only pace but potentially restart stalled recovery sequences, transforming our mechanistic insight into a foundation for novel therapeutic strategies aimed at restoring conscious awareness.

In conclusion, this study demonstrates that consciousness recovery from anesthesia is a hierarchical, dynamically reconfigured process actively orchestrated by CM. The multiscale framework we introduce quantifies this process, anchoring theoretical predictions in specific neural mechanisms and dynamic brain-wide signature. By moving beyond a binary switch modal to a detailed cascade, our work paves the way for predicting, monitoring, and ultimately facilitating the restoration of consciousness through objective biomarker-based monitoring and CM-targeted neuromodulation.

### Limitations of the study

Although our behavioral paradigms (DRVT and MRDR) were carefully designed to probe the restoration of consciousness level and content in mice, they remain inherently limited in capturing self-referential aspects of awareness. In addition, the awake, head-fixed fMRI approach, though enabling dynamic whole-brain mapping, may still introduce confounds related to restraint and reduced behavioral freedom, despite extensive animal habituation. Finally, while heart-rate stabilization correlated strongly with consciousness recovery in mice and aligned with retrospective clinical trends, further prospective validation in diverse patient populations is required to establish its reliability as a standalone biomarker.

## RESOURCE AVAILABILITY

### Lead contact

Further information and requests for resources and reagents should be directed to and will be fulfilled by the lead contact, Feng Wang.

### Materials availability

This study did not generate new unique reagents.

### Data and code availability

- Source data for quantitative comparisons presented herein will be deposited at Zenodo (https://doi.org/10.5281/zenodo.18501123) and made publicly available as of the date of publication.
- Codes to reproduce quantitative comparisons and visualization presented herein will be deposited at Zenodo (https://doi.org/10.5281/zenodo.18501123) and made publicly available as of the date of publication
- Any additional information required to reanalyze the data reported in this paper is available from the lead contact upon request.

## ACKNOWLEDGMENTS

We would like to express our gratitude to Haoran Liu for designing the behavioral box, as well as Xiang Li, Xulin Li, Gaoyang Zhao, Jialin Ye, Kang Huang, Qin Yang, Yaning Han, Yuting Tseng, and Binghao Zhao for their technical assistance. We extend our special thanks to MOVER for generously providing the extensive raw medical dataset. We also thank Helmut Kettenmann and Tianwen Huang for their valuable comments and feedback on the manuscript. This work was supported by the Brain Science and Brain-like Intelligence Technology-National Science and Technology Major Project (2022ZD0208300 to F.W.), the Shenzhen Medical Research Fund (B2502030 to F.W.), the National Natural Science Foundation of China (32371062 to F.W., 32230042 and 31930047 to L.W.), Financial Support for Outstanding Talents Training Fund in Shenzhen (to L.W.), the Guangdong Provincial Key Laboratory of Brain Connectome and Behavior (2023B1212060055 to L.W.), and the Scientific Instrument Developing Project of the Chinese Academy of Sciences (YJKYYQ20170064).

## AUTHOR CONTRIBUTIONS

F.W., and L.W. designed the project and proposed the concept. F.W., Y.Z., L.W., and S.H. wrote the manuscript with input from all authors. Y.Z., X.W., Y.X., and Y.T. performed behavior experiments and analyzed data. Y.Z. performed surgeries and calcium imaging experiments. Y.Z. and X.W. performed opto- and chemo-genetic manipulations and morphological analysis. Y.Z., Y.Y., and H.W. performed the multimodal data analysis. Y.Z. analyzed human data. Y.Z. designed the head-fixed awake fMRI scanning setup, performed, and analyzed the fMRI data. Y.Z., X.W., Y.T., and Y.Y. visualized the data.

## DECLARATION OF GENERATIVE AI AND AI-ASSISTED TECHNOLOGIES IN THE WRITING PROCESS

During the preparation of this work, the AI tool DeepSeek and Monica were used to assist with language editing. Upon its use, the authors thoroughly reviewed and further edited the entire text. The authors take full responsibility for the content of the publication.

## STAR ★ METHODS

Detailed methods are provided in the online version of this paper and include the following:

### KEY RESOURCES TABLE

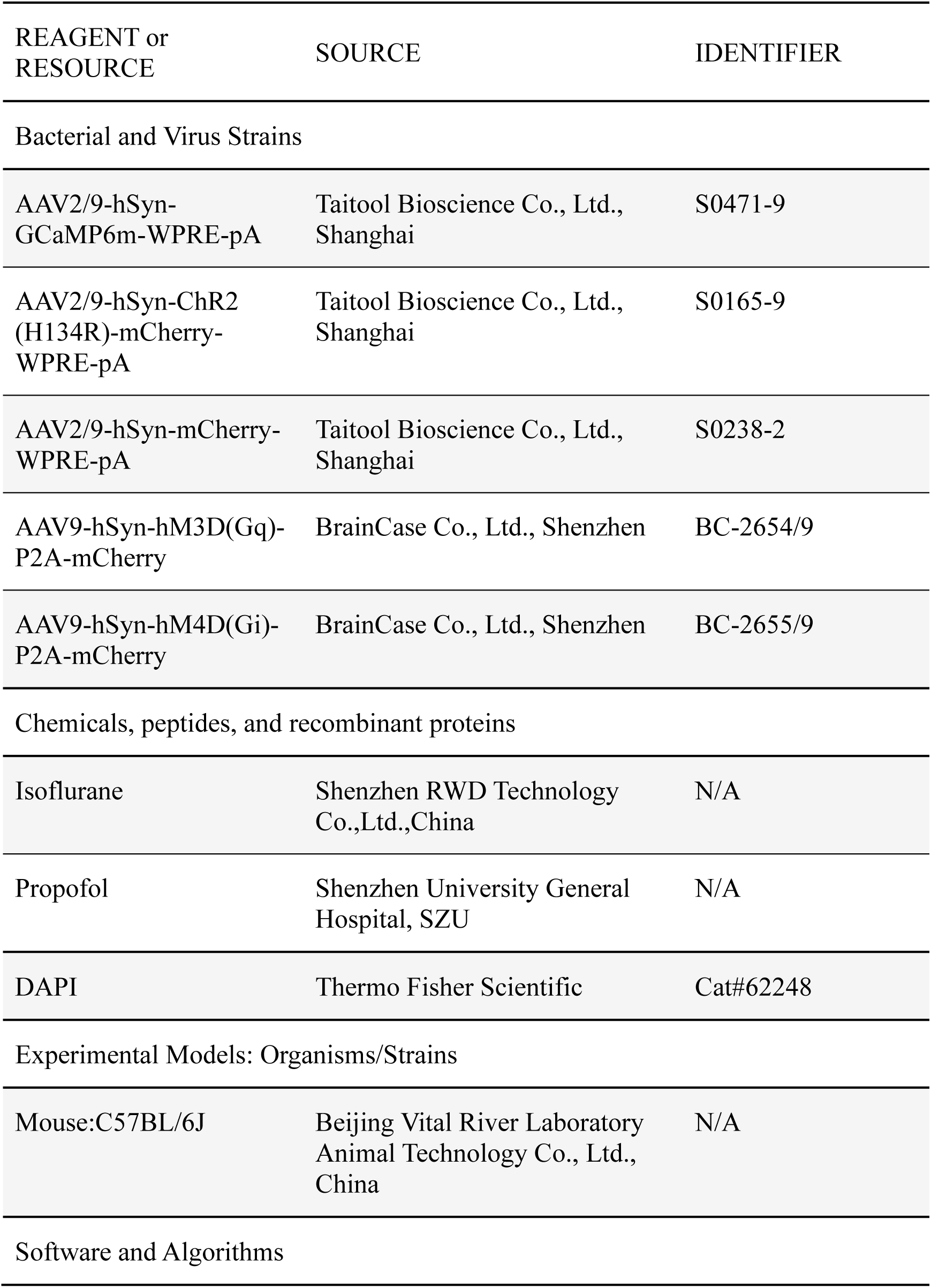
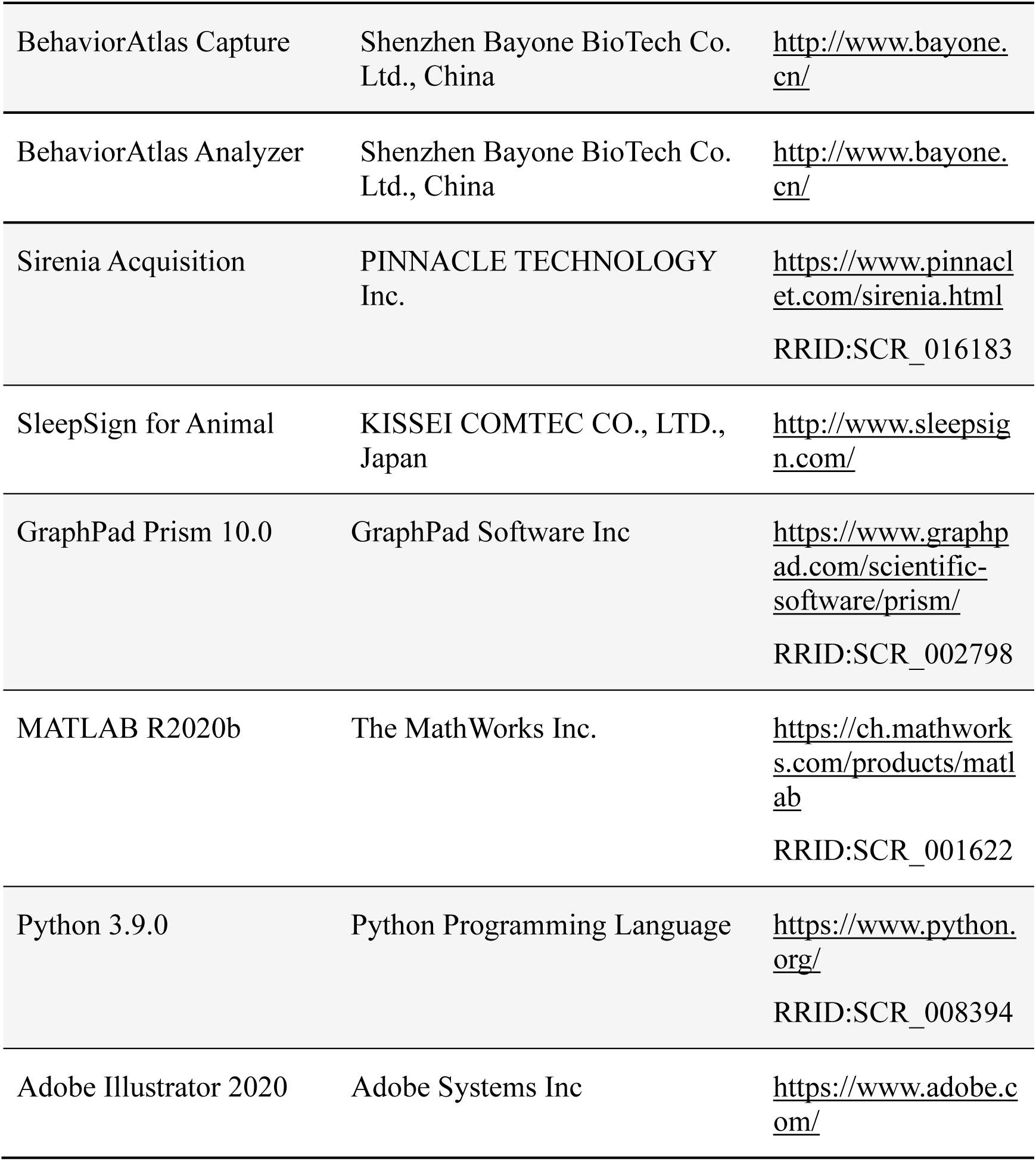

## EXPERIMENTAL MODEL AND STUDY PARTICIPANT DETAILS

### Animals

Adult male and female C57BL/6J mice (8-14 weeks old, Beijing Vital River Laboratory Animal Technology Co., Ltd., China) were used as experimental subjects. The mice were group-housed with 4-5 animals per cage, except when single-housing was required for specific experimental procedures. They were housed under controlled conditions with ad libitum access to food and water, a temperature range of 22-25°C, relative humidity of 30-70%, and a 12-h light-dark cycle (lights on at 08:00). Behavioral experiments were conducted daily during the same period (08:00-15:00). All animal husbandry and experimental procedures were conducted in accordance with guidelines approved by the Animal Care and Use Committee at the Shenzhen Institutes of Advanced Technology, Chinese Academy of Sciences, China (Approval number SIAT-IACUC-20210226-ZKYSZXJJSYJY-RZC-WF-A0923-03) and the Shenzhen Brain Science Infrastructure (Approval number SIAT-BSI-IRB-240329-NS-WF-A0001).

## METHOD DETAILS

### General anesthesia procedure

As described previously,^22,58^ general anesthesia was induced with 4% (vol/vol) isoflurane (RWD, China) in 100% oxygen. After the loss of righting reflex (LORR)^59^ was observed, general anesthesia was maintained with 1.4% isoflurane in 100% oxygen at a flow rate of 1.5 L/min for 30 minutes within the anesthesia induction chamber. For comparation, a separate group of mice received an intraperitoneal injection of propofol (100 mg/kg, Shenzhen University General Hospital, China). Once the anesthetics were discontinued, mice exhibiting LORR were carefully ang promptly moved to the behavioral testing apparatuses.

### Implantation of DSI radio-telemetric transmitters

Telemetry electrocardiogram (ECG), blood pressure (BP), and pulse rate metrics were utilized in the cardiovascular analyses. Mice were implanted with a DSI radio-telemetric transmitter (model HD-X11; DSI, St. Paul, MN, USA) for telemetry ECG-BP recording. The surgical procedure for implanting combined telemetric transmitters was performed as described previously.^60^ Briefly, mice were anesthetized with pentobarbital (100 mg/kg, i.p.). After shaving and disinfection of the cervical region, a midline skin incision (1-1.5 cm) was made below the chin. The left common carotid artery was isolated via blunt dissection, meticulously avoiding the vagus nerve. The pressure catheter integrated with the telemetry transmitter was introduced into the artery and advanced to position its tip within the aortic arch, then secured with three sutures. A subcutaneous pocket was created on the left flank using blunt dissection, irrigated with warm sterile saline, and the transmitter body was implanted. ECG leads were trimmed to appropriate lengths, stripped at the tips, and placed in Einthoven II configuration: the negative lead fixed to the right pectoral muscle and the positive lead looped and secured at the left caudal rib region using absorbable sutures, ensuring leads lay flat. The skin incision was closed with non-absorbable sutures and tissue adhesive. To prevent postoperative pain and infection, the surgical incision was treated with lincomycin-lidocaine gel. The mice were housed individually and administered meloxicam (0.5 mg/kg) daily for a 7-day recovery period before telemetric recording.

### Integrated 3D behavioral and cardiovascular recording

We established a synchronous multimodal acquisition platform by integrating a multi-view video capture device (described previously^61,62^) with implantable DSI telemetry (Matrix 2.0). This configuration enabled simultaneous monitoring of spontaneous and defensive behaviors alongside cardiovascular parameters in mice implanted with the combined ECG-BP transmitters. Behavioral data were captured at 30 Hz (640 × 480 pixels) using four Intel RealSense D435 cameras, while cardiovascular signals were recorded at 1 kHz (ECG) and 500 Hz (BP).

Spontaneous behavior test conducted in a transparent acrylic chamber (40 × 40 × 30 cm) under illumination from an overhead 27-inch LCD monitor. Mice underwent 60-minute recording sessions following 10-minute habituation 24 hours prior to testing. Data were collected during conscious baseline and throughout post-anesthesia recovery (initiated immediately upon anesthetic discontinuation). Defensive behavior paradigms utilized chambers (40 × 40 × 30 cm) connected to refuge compartments (15 × 18 × 15 cm for each chamber) with overhead looming stimuli (2°–20° expanding disc, described previously^61,63^) presented on the LCD monitor. Three chamber configurations were implemented: (1) single-refuge for DRVT in cue-unitary environments, (2) ipsilateral true-fake refuge (ips T-F) containing one functional and one non-functional refuge; and (3) multi-refuge configuration (1T-3F) with one functional and three non-functional refuges for assessing MRDR in cue-complex environments. Functional refuge permitted physical entry through an access door, while non-functional refuges were visually identical but physically inaccessible. All refuges units shared identical geometry, dimensions, and material properties (transparent acrylic with white plastic base). Following 10-minute habituation one day before testing, DRVT and MRDR were quantified during conscious baseline and at designated post-anesthesia timepoints (PR0, PR5, PR10, PR15).

### Multimodal behavioral and cardiovascular analysis

Behavioral (fps 30) and cardiovascular data were temporally aligned through frame-rate conversion, with cardiovascular records synchronized to behavioral frames using the formula:

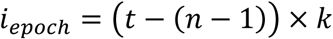

where *t* represents behavioral time in seconds, *n* denotes integer seconds, and *k* indicates the sampling density of cardiovascular data per second, achieving millisecond-level synchronization precision. Cardiovascular outliers were filtered using a15-frame rolling median correction with parameter-specific range validation, followed by forward-backward missing value imputation.

For spontaneous behavioral analysis, data were segmented into uniform 60-second intervals (1,800 frames) to capture periodic cardiovascular rhythms synchronized with behavioral patterns. Each epoch was characterized by: (1) cardiovascular dynamics (mean systolic/diastolic/mean pressure, heart rate variability (RMSSD), and first-derivative change rates); (2) kinematic metrics (velocity profiles, body curvature, and 3D spatial dispersion); and (3) behavioral complexity (Shannon entropy of movement states and Markov transition probabilities between behavioral categories). High-dimensional features were projected into 2D space using Uniform Manifold Approximation and Projection (UMAP) to visualize group separation (WF controls vs. PA cohorts) and temporal trajectories.

To quantify the dynamic recovery of consciousness, two complementary metrics were calculated: (1) feature-space divergence measured instantaneous deviation of PA from baseline consciousness (WF group):

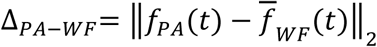

where Δ_*PA*−*WF*_ indicates scalar divergence, *f*_*PA*_(*t*) represents the feature vector of the PA group at time *t*, and *f*_*WF*_(*t*) is the time-matched WF baseline mean; and (2) normalized recovery index represented relative consciousness recovery on a 0-1 scale:

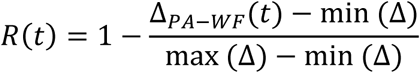

where *R*(*t*) indicates the normalized index, and min (Δ) and max (Δ) represent minimum and maximum divergence across all timepoints, respectively. Full consciousness recovery was defined as *R(t)* ≥ 0.95.

To identify the recovery stage, an integrated feature set encompassing autonomic regulation (normalized LF/HF powers, NN interval), cardiovascular function (mean HR, HR standard deviation), and behavioral state (frequency of movement transitions) was standardized across all intervals. Spectral clustering (clusters = 4, affinity = ‘nearest_neighbors’, neighbors = 10) was applied to the standardized 6-dimensional feature vectors from PA intervals to partition them into distinct recovery stages. Temporal distribution of each stage was characterized using kernel density estimation. Stage divergence from normal consciousness was quantified as the mean Euclidean distance from each PA epoch to its k-nearest neighbors (k = 5) within the WF baseline cohort. For defensive behavior clustering, a distinct 12-dimensional feature vector integrated flight behavior metrics (fraction, frequency), hemodynamic parameters (mean and maximum slopes of SysBP, DiaBP, MBP, and HR), and autonomic function (LF/HF ratio, NN interval). Following z-score normalization, features underwent UMAP dimensionality reduction and k-means clustering (k = 5, init = 10). Cluster segregation was illustrated using covariance-derived ellipses scaled to 2.5 standard deviations.

### fMRI acquisition and habituation

A custom-made 3D-printed headpost^64,65^ was surgically implanted under pentobarbital anesthesia (100 mg/kg, i.p.) for awake fMRI acquisition. Following scalp reflection and skull drying, the headpost was centrally positioned and secured using tissue glue adhesive (3M Vetbond^TM^) and dental cement, ensuring complete coverage of exposed tissue. Mice recovered for 7 days post-surgery before habituation.

A 14-day stepwised habituation protocol (Fig. 4a) acclimated mice to scanning conditions in a sound-attenuated chamber. Mice were head-fixed in an MRI-compatible cradle with body restraint using a concave arch. The protocol progressed as follows: days 1-2 (0.5 h fixation); days 3-4 (1 h); days 5-6 (1 h with 60 dB scanning noise); days 7-8 (1.5 h with 60 dB scanning noise); days 9-10 (1.5 with 90 dB scanning noise); day 11-12 (2 h with 90 dB noise); and days 13-14 (2 h with 110 dB noise). This protocol effectively minimizes motion and stress during scanning.

Functional imaging was performed on a 9.4 T scanner, 30-cm-diameter-bore magnet (uMR 9.4 T, United Imaging Life Science Instrument, P.R. China) equipped with a gradient insert with maximum strength at 1000 mT/m and a slew rate up to 10000 T/m/s. An 86 mm volume coil was used for transmission, and a 12 x 18 mm single-loop mouse head coil was used for reception. Each mouse was scanned sequentially across three states: awake baseline (WF), deeply anesthesia (1.4 % isoflurane, respiration 70-90 bpm), and post-anesthesia (PA) recovery. A T2 weighted FSE anatomical image (TR = 3000 ms, TE = 49.14 ms, matrix size = 331 × 368; FOV = 18 × 20 mm^2^; slice thickness = 500 μm; resolution = 50 × 50 μm^2^) was acquired for coregistration. Functional images were acquired using single-shot echo planar imaging (EPI, TR = 2000 ms, TE = 10.7 ms, FA = 60°, matrix size = 71 × 96, resolution = 200 × 200 μm^2^, slice thickness = 300 μm, slices = 54, volumes = 400).

### fMRI processing and dynamic functional connectivity

BOLD images were preprocessed in SPM12 and MATLAB 2020b.^64^ Steps included: manual brain extraction via ITK-SNAP; slice-timing and motion correction; spatial normalization (co-registration to subject-specific T2 anatomy, then nonlinear warping to the Allen Mouse Brain Atlas); and spatially smoothing (0.3 mm FWHM). BOLD signals were further processed by HRF-deconvolved and bandpass-filtered (0.01-0.1 Hz).

Dynamic functional connectivity (FC) was analyzed via a sliding window approach^66^ (window=30 TRs, step=10 TRs). Regional time series were z-scored, and dynamic FC matrices were clustered via k-means (k=3 optimized by silhouette scores) to identify recurring brain states. Graphic metrics (degree centrality, betweenness centrality, participation coefficient, global efficiency, modularity) were computed after retaining the top 10% of connections. Brain regions were assigned to 12 major functional networks^67^ (DMN, SMN, VIS, AUD, LIMB, TH, etc.; Table S5). Intranetwork and internetwork connectivity strengths were computed per state. Hub regions were identified by integrating autocorrelation changes, connectivity strength, and topological metrics via PCA-weighted multimodal scoring. Bootstrap stability analysis (100 iterations) validated hub consistency. Temporal staging of recovery was determined using Pruned Exact Linear Time change point detection applied to multi-dimensional features (state transitions, network properties, information-theoretic metrics). Stage-specific network configurations were derived by averaging FC matrices within each stage. The stage-specific causal strength of hub toward target regions was assessed by computing stage-averaged Granger causality. Statistical validation included Kruskal-Wallis tests for inter-state differences, permutation testing (1,000 iterations) against WF baselines, and FDR correction. Conscious baseline connectivity was defined as the mean of 400 preprocessed WF volumes. Visualizations (FC matrices, spatiotemporal heatmaps, network topology graphs, and Sankey diagrams) were generated in Python 3.9 using Nilearn, Scikit-learn, NetworkX, and Plotly libraries.

### Stereotactic injections and cannula implantation

Mice were anesthetized with pentobarbital (100 mg/kg, i.p.) and positioned on a stereotaxic apparatus (RWD, China) under continuous 1.5 % isoflurane maintenance. Ophthalmic ointment was applied to prevent corneal drying, followed by aseptic preparation of the surgical site. The scalp and connective tissues were removed to expose the skull. Stereotaxic coordinates relative to Bregma were as follows: Pef (anteroposterior (AP) −1.8 mm, mediolateral (ML) 1.0 mm, dorsoventral (DV) −5.1 mm), CM (AP −1.35 mm, ML 0 mm, DV −3.70 mm), and SC (AP −3.80 mm, ML 0.8 mm, DV −1.8 mm). Viral vectors were infused using a microsyringe pump (KDS Legato 130, RWD) connected to a 33-Ga needle (Hamilton) at 30 nl/min. For chemogenetic manipulation, AAV9-hSyn-hM4D(Gi)-P2A-mCherry (5.10 x 1012 genome copies per ml (gc/ml), BrainCase Co., Ltd., Shenzhen) or AAV9-hSyn-hM3D(Gq)-P2A-mCherry (5.06 x 1012 gc/ml, BrainCase) were unilaterally infused into CM (60 nl). For optogenetic manipulation, AAV2/9-hSyn-ChR2 (H134R)-mCherry-WPRE-pA (1.69 x 1013 gc/ml, Taitool Bioscience Co., Ltd., Shanghai) or AAV2/9-hSyn-mCherry-WPRE-pA (1.14 x 1013 gc/ml, Taitool) were infused into Pef (50 nl), CM (60nl), and SC (150nl). For fiber photometry, AAV2/9-hSyn-GCaMP6m-WPRE-pA (1.10 x 1013 gc/ml, Taitool) was infused into Pef (50 nl), CM (60 nl), and SC (150 nl). Following 3 weeks of viral expression, an optical fiber cannula (200-μm core, numerical aperture 0.37, Newdoon, China) was implanted at CM (AP −1.35 mm, ML 0 mm, DV: −3.50 mm), Pef (AP −1.8 mm, ML 1.0 mm, DV −4.9 mm), or SC (AP −3.80 mm, ML 0.8 mm, DV −1.6 mm), and affixed to the skull with dental cement. Mice were allowed to recover for 7-10 days post-surgery before further tests.

### Pinch test

Noxious mechanical stimuli were administered to mice as previously described.^68^ Each mouse was placed in a plexiglass cage, and an alligator clip (Generic Micro Steel Toothless Alligator Clips, 5AMP) generating 340 g of force was applied to the ventral skin surface between the footpad and heel. Behavioral responses during wakefulness and post-anesthesia periods were recorded using a digital video camera. The duration of licking and flinching behaviors within 60 seconds after stimuli were analyzed using Aegisub (version 3.4.2).

### Rotarod test

As previously described,^69^ mice were habituated on the stationary rod for 2 minutes prior to training. Training sessions were conducted over two consecutive days, with each session comprising 3 trials (5-minute inter-trial intervals). During trials, the rod accelerated to 20 m/min after fully conscious mice stabilized, maintaining rotation for 2 minutes. For testing, mice were placed on the rod during wakefulness, as well as at PR0, PR5, PR10, and PR15 following general anesthesia procedure. Each test trial used identical rotation parameters (20 m/min, 2-minute duration), with fall latency from the rod within trials recorded via a digital video camera and compared across periods.

### Treadmill test

The treadmill apparatus comprised six individual lanes equipped with electric shock grids. Continuous foot shocks were delivered when mice failed to maintain the preset speed and entered the electric shock grid. Prior to testing, mice underwent three consecutive days of training: each session consisted of 2 minutes of running at speed of 10 m/min with moderate shocks. If a mouse exhibited exhaustion, shocks were deactivated immediately, and the mouse was allowed to rest for 5 minutes before retraining. During formal testing, locomotor performance was assessed for 2 minutes during wakefulness, as well as at PR0, PR5, PR10 and PR15 following general anesthesia procedure. The cumulative shock numbers per trial were quantified and compared across stages.

### EEG/EMG recording and analysis

Stainless steel screw electrodes (Pinnacle, catalog #8209) were inserted into the skull^70^: two located 1.5 mm from the midline and 1.5 mm anterior to the bregma, and another two located 3 mm from the midline and 3.5 mm posterior to the bregma. Two EMG electrodes were inserted into neck musculature. Insulated leads from the EEG/EMG electrodes were soldered to a pin header, which was secured to the skull using dental cement. Following 7 days of postoperative recovery and 2 days of habituation, EEG/EMG data was collected using EEG/EMG recording system (8400-K1, PINNACLE) with a Pinnacle’s high-gain preamplifiers (X100 amplification, high-pass filter 0.5 Hz for EEG and 10 Hz for EMG, sampling rate, 2,000 Hz).

Raw EEG/EMG data were converted into European data format (.edf) and processed via a standardized Python 3.9 pipeline (MNE-Python v1.4). Spectral analysis employed Welch’s method (window: 4 s, overlap: 50%) to compute power spectral density (PSD) in frequency bands: delta (0.5-4 Hz), theta (4-8 Hz), alpha (8-13 Hz), beta (13-30 Hz) for EEG, and broadband EMG (10-300 Hz). Relative bandpower was derived by normalizing band-specific power against total power. Time-frequency representations (TFRs) were generated using Morlet wavelets (frequencies: 0.5-30 Hz for EEG, 10-300 Hz for EMG; cycles = freqs/2). Dynamic bandpower fluctuations were quantified via short-time Fourier transform (STFT; nperseg = 4×sfreq). Computational efficiency was optimized via downsampling (waveforms: 20 Hz) for batch processing.

### Pupillometry

A custom-made stainless metal head-plate was cemented to the skull using dental adhesive to stabilize an eye-tracking unit (ETU) for freely moving recordings.^71^ Following 7 days of postoperative recovery, pupil dynamics were monitored at 200 Hz using a head-mounted miniature infrared camera controlled by LabVIEW (2018, NI). Mice underwent 10-minute habituation in the recording chamber 24 hours prior to testing. A convolutional neural network trained via DeepLabCut (v2.1.8.2) by labeling 9 landmarks (8 boundary points and 1 center) was utilized to quantify pupil diameter. Pupil radius was computed as the mean Euclidean distance from boundary points to the center, with diameter derived as twice the radius. To quantify looming stimulus-evoked pupil dynamics, the mean pupil diameter during the 5 s preceding stimulus onset was defined as the baseline. This baseline value was compared to the mean diameter over the 30 s post-stimulus interval to calculate the percentage change in pupil size.

### Fiber photometry

Fluorescence signals were acquired using a dual-color fiber photometry system (NEWDOON) with 470-nm (GCaMP6m excitation) and 410-nm (isosbestic control) LEDs pulsed alternately at 100 Hz. Excitation light passed through bandpass filters (470±20 nm, 410±20 nm; CHROMA), while emission was collected via a 525±22.5 nm filter (SEMROCK) and detected by a photomultiplier tube (2 kHz sampling; Hamamatsu H10722-210). Signals were downsampled to 200 Hz and synchronized with behavior via system software. For analysis, raw data were smoothed (10-point moving average). Δ*F*/*F* was calculated by scaling the 410-nm trace to fit the 470-nm signal using least-squares regression. Activity was quantified via state-specific *ΔF*/*F* and areas under curve (AUC).

### Optogenetics and chemogenetics manipulations

Mice connected to an optic fiber and allowed to habituate in behavior box for 10 minutes one day prior to experimentation. During testing, mice were quickly transferred to a looming box immediately following isoflurane cessation while in a LORR state. A 5-s blue laser stimulus (473 nm) was manually delivered through the optic fiber, comprising 50 pulses at 20 Hz (5-ms pulse width, 8-10 mW power at fiber tip).^72^ Stimulus onset occurred within 5 s of placement in the behavioral arena to ensure neural modulation during PA period. For chemogenetics manipulation, mice were injected intraperitoneally with CNO (2 mg/kg, Clozapine N-oxide, C0832, Sigma, dissolved in saline mixed with DMSO) or vehicle (saline with DMSO) 30 minutes before start of general anesthesia procedure (lasting for 30 minutes). Following anesthetic discontinuation, mice exhibiting LORR were immediately and gently transferred to behavioral box.

### Histology and microscopy

After behavioral tests, mice were overdosed with sodium pentobarbital (100 mg/kg, i.p.) and transcardially perfused with PBS (pH 7.4) followed by 4% PFA (Boster) at 5 mL/min. Brains were post-fixed in 4% PFA at 4°C for 24 h, then cryoprotected in 30% sucrose at 4°C until sinking for 48 h. Coronal sections (30 μm) were cut on a cryostat microtome (Leica CM1950). The sections were washed with PBS and incubated with DAPI (Sigma) for 5 minutes. All sections were photographed using a virtual slide scanning system (Olympus VS120) and confocal microscope (Zeiss LSM 880).

## QUANTIFICATION AND STATISTICAL ANALYSIS

### Statistical analysis

The number of biological replicates in each group varied depending on the experiment and was indicated in figure legends. Before formal analysis, the normality of all datasets was assessed using the Shapiro-Wilk test (α=0.05), and homogeneity of variances was evaluated using Levene’s test. For normally distributed data, two-group comparisons used two-tailed *t*-tests, while multi-group comparisons employed one-way ANOVA with Dunnett’s post hoc test or two-way ANOVA with Bonferroni post-hoc test. Non-normally distributed data were analyzed using multiple Mann-Whitney tests with Bonferroni–Dunn correction for multiple comparisons (two groups), or using Kruskal-Wallis test, followed by Dunn’s post-hoc test with a Bonferroni correction for multiple pairwise comparisons (≥3 groups). All analyses were performed in GraphPad Prism 10.0, with significance thresholds for adjusted *P* values set at \**P*<0.05, \*\**P<0.01*, \*\*\**P*<0.001; exact *P* values and test parameters are provided in supplementary tables. Pooled data from male and female mice were presented as mean, mean ± sem, median with 95% bootstrap confidence intervals, or boxplots (medians (center line), IQR (box bounds), 1.5×IQR whiskers, outliers as individual points).

## REFERENCES

1. Hameroff, Stuart R. (2006). The Entwined Mysteries of Anesthesia and Consciousness: Is There a Common Underlying Mechanism? Anesthesiology 105, 400–412. 10.1097/00000542-200608000-00024.

2. Mashour, G.A. (2024). Anesthesia and the neurobiology of consciousness. Neuron 112, 1553–1567. 10.1016/j.neuron.2024.03.002.

3. Mudrik, L., Boly, M., Dehaene, S., Fleming, S.M., Lamme, V., Seth, A., and Melloni, L. (2025). Unpacking the complexities of consciousness: Theories and reflections. Neurosci Biobehav Rev 170, 106053. 10.1016/j.neubiorev.2025.106053.

4. Mashour, G.A., Roelfsema, P., Changeux, J.P., and Dehaene, S. (2020). Conscious Processing and the Global Neuronal Workspace Hypothesis. Neuron 105, 776–798. 10.1016/j.neuron.2020.01.026.

5. Tononi, G., Boly, M., Massimini, M., and Koch, C. (2016). Integrated information theory: from consciousness to its physical substrate. Nature Reviews Neuroscience 17, 450–461. 10.1038/nrn.2016.44.

6. Seth, A.K., and Bayne, T. (2022). Theories of consciousness. Nat Rev Neurosci 23, 439–452. 10.1038/s41583-022-00587-4.

7. Alkire, M.T., Hudetz, A.G., and Tononi, G. (2008). Consciousness and Anesthesia. Science 322, 876–880. 10.1126/science.1149213.

8. Mashour, G.A., Palanca, B.J., Basner, M., Li, D., Wang, W., Blain-Moraes, S., Lin, N., Maier, K., Muench, M., Tarnal, V., Vanini, G., Ochroch, E.A., Hogg, R., Schwartz, M., Maybrier, H., Hardie, R., Janke, E., Golmirzaie, G., Picton, P., McKinstry-Wu, A.R., Avidan, M.S., and Kelz, M.B. (2021). Recovery of consciousness and cognition after general anesthesia in humans. Elife 10. 10.7554/eLife.59525.

9. Mudrik, L., Hirschhorn, R., and Korisky, U. (2024). Taking consciousness for real: Increasing the ecological validity of the study of conscious vs. unconscious processes. Neuron 112, 1642–1656. 10.1016/j.neuron.2024.03.031.

10. Kronemer, S.I., Bandettini, P.A., and Gonzalez-Castillo, J. (2025). Sleuthing subjectivity: a review of covert measures of consciousness. Nat Rev Neurosci 26, 476–496. 10.1038/s41583-025-00934-1.

11. Brown, E.N., Lydic, R., and Schiff, N.D. (2010). General anesthesia, sleep, and coma. N Engl J Med 363, 2638–2650. 10.1056/NEJMra0808281.

12. Robba, C., Graziano, F., Rebora, P., Elli, F., Giussani, C., Oddo, M., Meyfroidt, G., Helbok, R., Taccone, F.S., Prisco, L., Vincent, J.L., Suarez, J.I., Stocchetti, N., and Citerio, G. (2021). Intracranial pressure monitoring in patients with acute brain injury in the intensive care unit (SYNAPSE-ICU): an international, prospective observational cohort study. Lancet Neurol 20, 548–558. 10.1016/s1474-4422(21)00138-1.

13. Katoh, T., Ikeda, K., and Bito, H. (1997). Does nitrous oxide antagonize sevoflurane-induced hypnosis? Br J Anaesth 79, 465–468. 10.1093/bja/79.4.465.

14. Bodien, Y.G., Vora, I., Barra, A., Chiang, K., Chatelle, C., Goostrey, K., Martens, G., Malone, C., Mello, J., Parlman, K., Ranford, J., Sterling, A., Waters, A.B., Hirschberg, R., Katz, D.I., Mazwi, N., Ni, P., Velmahos, G., Waak, K., Edlow, B.L., and Giacino, J.T. (2023). Feasibility and Validity of the Coma Recovery Scale-Revised for Accelerated Standardized Testing: A Practical Assessment Tool for Detecting Consciousness in the Intensive Care Unit. Ann Neurol 94, 919–924. 10.1002/ana.26740.

15. Mehta, R., and Chinthapalli, K. (2019). Glasgow coma scale explained. Bmj 365, l1296. 10.1136/bmj.l1296.

16. Brennan, P.M., Whittingham, C., Sinha, V.D., and Teasdale, G. (2024). Assessment of level of consciousness using Glasgow Coma Scale tools. Bmj 384, e077538. 10.1136/bmj-2023-077538.

17. Makarem, J., Larijani, A.H., Eslami, B., Jafarzadeh, A., Karvandian, K., and Mireskandari, S.M. (2020). Risk factors of inadequate emergence following general anesthesia with an emphasis on patients with substance dependence history. Korean J Anesthesiol 73, 302–310. 10.4097/kja.19214.

18. Sumner, M., Deng, C., Evered, L., Frampton, C., Leslie, K., Short, T., and Campbell, D. (2023). Processed electroencephalography-guided general anaesthesia to reduce postoperative delirium: a systematic review and meta-analysis. British Journal of Anaesthesia 130, e243–e253. 10.1016/j.bja.2022.01.006.

19. Fischer, D., Edlow, B.L., Freeman, H.J., Alaiev, D., Wu, Q., Ware, J.B., Detre, J.A., and Aguirre, G.K. (2025). Reconstructing Covert Consciousness: Neural Decoding as a Novel Consciousness Assessment. Neurology 104, e210208. 10.1212/wnl.0000000000210208.

20. Ye, J., Xu, Y., Huang, K., Wang, X., Wang, L., and Wang, F. (2025). Hierarchical behavioral analysis framework as a platform for standardized quantitative identification of behaviors. Cell Rep 44, 115239. 10.1016/j.celrep.2025.115239.

21. Coma, C.T. (2008). Dimensions of Consciousness: Nosology of Disorders of Consciousness Arousal and Awareness. Psychology.

22. Zhong, H., Tong, L., Gu, N., Gao, F., Lu, Y., Xie, R.G., Liu, J., Li, X., Bergeron, R., Pomeranz, L.E., Mackie, K., Wang, F., Luo, C.X., Ren, Y., Wu, S.X., Xie, Z., Xu, L., Li, J., Dong, H., Xiong, L., and Zhang, X. (2017). Endocannabinoid signaling in hypothalamic circuits regulates arousal from general anesthesia in mice. J Clin Invest 127, 2295–2309. 10.1172/jci91038.

23. Pfaff, D., Ribeiro, A., Matthews, J., and Kow, L.M. (2008). Concepts and mechanisms of generalized central nervous system arousal. Ann N Y Acad Sci 1129, 11–25. 10.1196/annals.1417.019.

24. Garey, J., Goodwillie, A., Frohlich, J., Morgan, M., Gustafsson, J.-A., Smithies, O., Korach, K.S., Ogawa, S., and Pfaff, D.W. (2003). Genetic contributions to generalized arousal of brain and behavior. Proceedings of the National Academy of Sciences 100, 11019–11022. doi:10.1073/pnas.1633773100.

25. Raut, R.V., Rosenthal, Z.P., Wang, X., Miao, H., Zhang, Z., Lee, J.-M., Raichle, M.E., Bauer, A.Q., Brunton, S.L., Brunton, B.W., and Kutz, J.N. (2025). Arousal as a universal embedding for spatiotemporal brain dynamics. Nature 647, 454–461. 10.1038/s41586-025-09544-4.

26. Samad, M., Angel, M., Rinehart, J., Kanomata, Y., Baldi, P., and Cannesson, M. (2023). Medical Informatics Operating Room Vitals and Events Repository (MOVER): a public-access operating room database. JAMIA Open 6, ooad084. 10.1093/jamiaopen/ooad084.

27. Song, X.J., and Hu, J.J. (2024). Neurobiological basis of emergence from anesthesia. Trends Neurosci 47, 355–366. 10.1016/j.tins.2024.02.006.

28. Yu, Q., Wang, Y., Gu, L., Shao, W., Gu, J., Liu, L., Lian, X., Xu, Q., Zhang, Y., Yang, Y., Zhang, Z., Wu, Y., Ma, H., Shen, Y., Ye, W., Wu, Y., Yang, H., Chen, L., Nagayasu, K., and Zhang, H. (2024). Dorsal raphe nucleus to basolateral amygdala 5-HTergic neural circuit modulates restoration of consciousness during sevoflurane anesthesia. Biomedicine & Pharmacotherapy 176, 116937. 10.1016/j.biopha.2024.116937.

29. Zhang, R., Gan, X., Xu, T., Yu, F., Wang, L., Song, X., Jiao, G., Liu, X., Zhou, F., and Becker, B. (2025). A neurofunctional signature of affective arousal generalizes across valence domains and distinguishes subjective experience from autonomic reactivity. Nature Communications 16, 6492. 10.1038/s41467-025-61706-0.

30. Min, B.K. (2010). A thalamic reticular networking model of consciousness. Theor Biol Med Model 7, 10. 10.1186/1742-4682-7-10.

31. Schiff, N.D. (2020). Central Lateral Thalamic Nucleus Stimulation Awakens Cortex via Modulation of Cross-Regional, Laminar-Specific Activity during General Anesthesia. Neuron 106, 1–3. 10.1016/j.neuron.2020.02.016.

32. Roy, D.S., Zhang, Y., Halassa, M.M., and Feng, G. (2022). Thalamic subnetworks as units of function. Nature Neuroscience 25, 140–153. 10.1038/s41593-021-00996-1.

33. Seth, A.K., Barrett, A.B., and Barnett, L. (2015). Granger causality analysis in neuroscience and neuroimaging. J Neurosci 35, 3293–3297. 10.1523/jneurosci.4399-14.2015.

34. Fang, Z., Dang, Y., Ping, A., Wang, C., Zhao, Q., Zhao, H., Li, X., and Zhang, M. (2025). Human high-order thalamic nuclei gate conscious perception through the thalamofrontal loop. Science 388, eadr3675. 10.1126/science.adr3675.

35. Liu, X., Huang, H., Snutch, T.P., Cao, P., Wang, L., and Wang, F. (2022). The Superior Colliculus: Cell Types, Connectivity, and Behavior. Neurosci Bull 38, 1519–1540. 10.1007/s12264-022-00858-1.

36. Hoy, J.L., and Farrow, K. (2025). The superior colliculus. Current Biology 35, R164–R168. 10.1016/j.cub.2025.01.022.

37. Kostin, A., Alam, M.A., Saevskiy, A., McGinty, D., and Alam, M.N. (2022). Activation of the Ventrolateral Preoptic Neurons Projecting to the Perifornical-Hypothalamic Area Promotes Sleep: DREADD Activation in Wild-Type Rats. Cells 11. 10.3390/cells11142140.

38. Huo, J., Li, H., Wang, D., Wang, S., Zhang, X., Dong, H., and Li, J. (2025). Orexin signalling in the nucleus accumbens promotes arousal from isoflurane anaesthesia and restores communication between the nucleus accumbens and frontal cortex. Br J Anaesth 135, 941–952. 10.1016/j.bja.2025.03.042.

39. Phillips, I. (2018). The methodological puzzle of phenomenal consciousness. Philos Trans R Soc Lond B Biol Sci 373. 10.1098/rstb.2017.0347.

40. Fang, Z., Dang, Y., Li, X., Zhao, Q., Zhang, M., and Zhao, H. (2024). Intracranial neural representation of phenomenal and access consciousness in the human brain. Neuroimage 297, 120699. 10.1016/j.neuroimage.2024.120699.

41. Haun, A.M. (2022). IIT is ideally positioned to explain perceptual phenomena. Behav Brain Sci 45, e52. 10.1017/s0140525x21001965.

42. Seth, A.K., and Friston, K.J. (2016). Active interoceptive inference and the emotional brain. Philos Trans R Soc Lond B Biol Sci 371. 10.1098/rstb.2016.0007.

43. Owens, A.P., Allen, M., Ondobaka, S., and Friston, K.J. (2018). Interoceptive inference: From computational neuroscience to clinic. Neurosci Biobehav Rev 90, 174–183. 10.1016/j.neubiorev.2018.04.017.

44. Critchley, H.D., Wiens, S., Rotshtein, P., Ohman, A., and Dolan, R.J. (2004). Neural systems supporting interoceptive awareness. Nat Neurosci 7, 189–195. 10.1038/nn1176.

45. Klincewicz, M., Cheng, T., Schmitz, M., Sebastián, M., and Snyder, J.S. (2025). What makes a theory of consciousness unscientific? Nat Neurosci 28, 689–693. 10.1038/s41593-025-01881-x.

46. Seth, A.K. (2013). Interoceptive inference, emotion, and the embodied self. Trends Cogn Sci 17, 565–573. 10.1016/j.tics.2013.09.007.

47. LeDoux, J.E., and Brown, R. (2017). A higher-order theory of emotional consciousness. Proc Natl Acad Sci U S A 114, E2016–E2025. 10.1073/pnas.1619316114.

48. Brown, R., Lau, H., and LeDoux, J.E. (2019). Understanding the Higher-Order Approach to Consciousness. Trends in Cognitive Sciences 23, 754–768. 10.1016/j.tics.2019.06.009.

49. Bachmann, T., Suzuki, M., and Aru, J. (2020). Dendritic integration theory: A thalamo-cortical theory of state and content of consciousness. Philosophy and the Mind Sciences 1. 10.33735/phimisci.2020.II.52.

50. Aru, J., Larkum, M., and Shine, J. (2023). The feasibility of artificial consciousness through the lens of neuroscience 10.48550/arXiv.2306.00915.

51. Cogitate, C., Ferrante, O., Gorska-Klimowska, U., Henin, S., Hirschhorn, R., Khalaf, A., Lepauvre, A., Liu, L., Richter, D., Vidal, Y., Bonacchi, N., Brown, T., Sripad, P., Armendariz, M., Bendtz, K., Ghafari, T., Hetenyi, D., Jeschke, J., Kozma, C., Mazumder, D.R., Montenegro, S., Seedat, A., Sharafeldin, A., Yang, S., Baillet, S., Chalmers, D.J., Cichy, R.M., Fallon, F., Panagiotaropoulos, T.I., Blumenfeld, H., de Lange, F.P., Devore, S., Jensen, O., Kreiman, G., Luo, H., Boly, M., Dehaene, S., Koch, C., Tononi, G., Pitts, M., Mudrik, L., and Melloni, L. (2025). Adversarial testing of global neuronal workspace and integrated information theories of consciousness. Nature 642, 133–142. 10.1038/s41586-025-08888-1.

52. Casali, A.G., Gosseries, O., Rosanova, M., Boly, M., Sarasso, S., Casali, K.R., Casarotto, S., Bruno, M.A., Laureys, S., Tononi, G., and Massimini, M. (2013). A theoretically based index of consciousness independent of sensory processing and behavior. Sci Transl Med 5, 198ra105. 10.1126/scitranslmed.3006294.

53. Melloni, L., Mudrik, L., Pitts, M., and Koch, C. (2021). Making the hard problem of consciousness easier. Science 372, 911–912. 10.1126/science.abj3259.

54. Yurchenko, S.B. (2024). Panpsychism and dualism in the science of consciousness. Neurosci Biobehav Rev 165, 105845. 10.1016/j.neubiorev.2024.105845.

55. Gao, S., and Calderon, D.P. (2020). Robust alternative to the righting reflex to assess arousal in rodents. Sci Rep 10, 20280. 10.1038/s41598-020-77162-3.

56. Vatner, S.F. (1978). Effects of anesthesia on cardiovascular control mechanisms. Environ Health Perspect 26, 193–206. 10.1289/ehp.7826193.

57. Tasserie, J., Uhrig, L., Sitt, J.D., Manasova, D., Dupont, M., Dehaene, S., and Jarraya, B. (2022). Deep brain stimulation of the thalamus restores signatures of consciousness in a nonhuman primate model. Sci Adv 8, eabl5547. 10.1126/sciadv.abl5547.

58. Haruwaka, K., Ying, Y., Liang, Y., Umpierre, A.D., Yi, M.H., Kremen, V., Chen, T., Xie, T., Qi, F., Zhao, S., Zheng, J., Liu, Y.U., Dong, H., Worrell, G.A., and Wu, L.J. (2024). Microglia enhance post-anesthesia neuronal activity by shielding inhibitory synapses. Nat Neurosci 27, 449–461. 10.1038/s41593-023-01537-8.

59. Leung, L.S., Luo, T., Ma, J., and Herrick, I. (2014). Brain areas that influence general anesthesia. Prog Neurobiol 122, 24–44. 10.1016/j.pneurobio.2014.08.001.

60. Au - Rötzer, R.D., Au - Brox, V.F., Au - Hennis, K., Au - Thalhammer, S.B., Au - Biel, M., Au - Wahl-Schott, C., and Au - Fenske, S. (2021). Implantation of Combined Telemetric ECG and Blood Pressure Transmitters to Determine Spontaneous Baroreflex Sensitivity in Conscious Mice. JoVE, e62101. doi:10.3791/62101.

61. Liu, X., Feng, X., Huang, H., Huang, K., Xu, Y., Ye, S., Tseng, Y.T., Wei, P., Wang, L., and Wang, F. (2022). Male and female mice display consistent lifelong ability to address potential life-threatening cues using different post-threat coping strategies. BMC Biol 20, 281. 10.1186/s12915-022-01486-x.

62. Huang, K., Han, Y., Chen, K., Pan, H., Zhao, G., Yi, W., Li, X., Liu, S., Wei, P., and Wang, L. (2021). A hierarchical 3D-motion learning framework for animal spontaneous behavior mapping. Nat Commun 12, 2784. 10.1038/s41467-021-22970-y.

63. Yilmaz, M., and Meister, M. (2013). Rapid innate defensive responses of mice to looming visual stimuli. Curr Biol 23, 2011–2015. 10.1016/j.cub.2013.08.015.

64. Yu, Y., Qiu, Y., Li, G., Zhang, K., Bo, B., Pei, M., Ye, J., Thompson, G.J., Cang, J., Fang, F., Feng, Y., Duan, X., Tong, C., and Liang, Z. (2023). Sleep fMRI with simultaneous electrophysiology at 9.4 T in male mice. Nature Communications 14, 1651. 10.1038/s41467-023-37352-9.

65. Gutierrez-Barragan, D., Singh, N.A., Alvino, F.G., Coletta, L., Rocchi, F., De Guzman, E., Galbusera, A., Uboldi, M., Panzeri, S., and Gozzi, A. (2022). Unique spatiotemporal fMRI dynamics in the awake mouse brain. Current Biology 32, 631–644.e636. 10.1016/j.cub.2021.12.015.

66. Huang, Z., Tarnal, V., Vlisides, P.E., Janke, E.L., McKinney, A.M., Picton, P., Mashour, G.A., and Hudetz, A.G. (2021). Asymmetric neural dynamics characterize loss and recovery of consciousness. Neuroimage 236, 118042. 10.1016/j.neuroimage.2021.118042.

67. Gutierrez-Barragan, D., Ramirez, J.S.B., Panzeri, S., Xu, T., and Gozzi, A. (2024). Evolutionarily conserved fMRI network dynamics in the mouse, macaque, and human brain. Nat Commun 15, 8518. 10.1038/s41467-024-52721-8.

68. Huang, T.W., Lin, S.H., Malewicz, N.M., Zhang, Y., Zhang, Y., Goulding, M., LaMotte, R.H., and Ma, Q.F. (2019). Identifying the pathways required for coping behaviours associated with sustained pain. Nature 565, 86-+. 10.1038/s41586-018-0793-8.

69. Shan, H.M., Maurer, M.A., and Schwab, M.E. (2023). Four-parameter analysis in modified Rotarod test for detecting minor motor deficits in mice. BMC Biol 21, 177. 10.1186/s12915-023-01679-y.

70. Tseng, Y.T., Zhao, B., Chen, S., Ye, J., Liu, J., Liang, L., Ding, H., Schaefke, B., Yang, Q., Wang, L., Wang, F., and Wang, L. (2022). The subthalamic corticotropin-releasing hormone neurons mediate adaptive REM-sleep responses to threat. Neuron 110, 1223–1239.e1228. 10.1016/j.neuron.2021.12.033.

71. Huang, K., Yang, Q., Han, Y., Zhang, Y., Wang, Z., Wang, L., and Wei, P. (2022). An Easily Compatible Eye-tracking System for Freely-moving Small Animals. Neurosci Bull 38, 661–676. 10.1007/s12264-022-00834-9.

72. Zhou, Z., Liu, X., Chen, S., Zhang, Z., Liu, Y., Montardy, Q., Tang, Y., Wei, P., Liu, N., Li, L., Song, R., Lai, J., He, X., Chen, C., Bi, G., Feng, G., Xu, F., and Wang, L. (2019). A VTA GABAergic Neural Circuit Mediates Visually Evoked Innate Defensive Responses. Neuron 103, 473–488.e476. 10.1016/j.neuron.2019.05.027.

