## Supplemental Figure 1∽11 for "Multimodal profiling reveals the centromedian nucleus of thalamus as a dynamic hub orchestrating staged consciousness recovery"

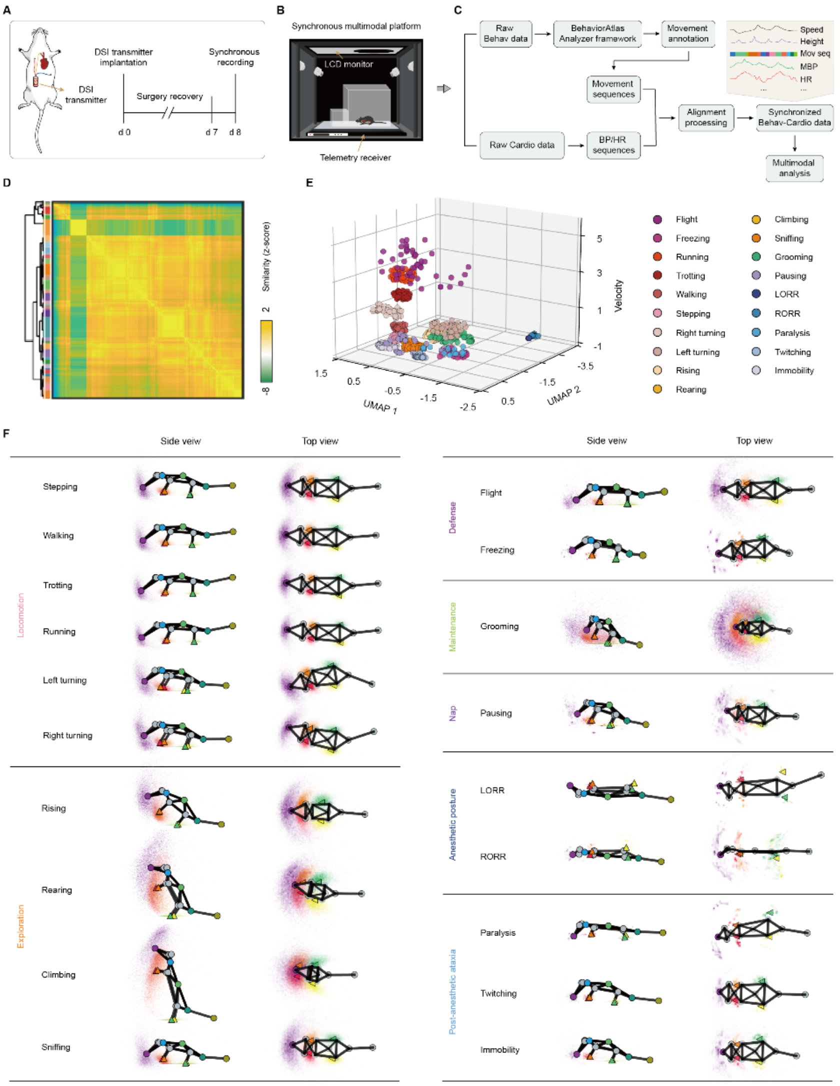


**Figure S1. An integrated multimodal framework for recording, analysis, and movement taxonomy, related to Figure 1**

(A) Paradigm of DSI transmitter implantation and timeline of synchronous behavioral-cardiovascular recording.

(B) Experimental setup for synchronous behavioral-cardiovascular monitoring.

(C) Multimodal data processing pipline.

(D) Movement similarity matrix (top) and its hierarchical clustering (bottom).

(E) UMAP visualization of movements categorized.

(F) Average movement skeletons (side and top views). Abbreviation: Behav, behavioral; Cardio, Cardiovascular; BP, Blood pressure; HR, Heart rate.

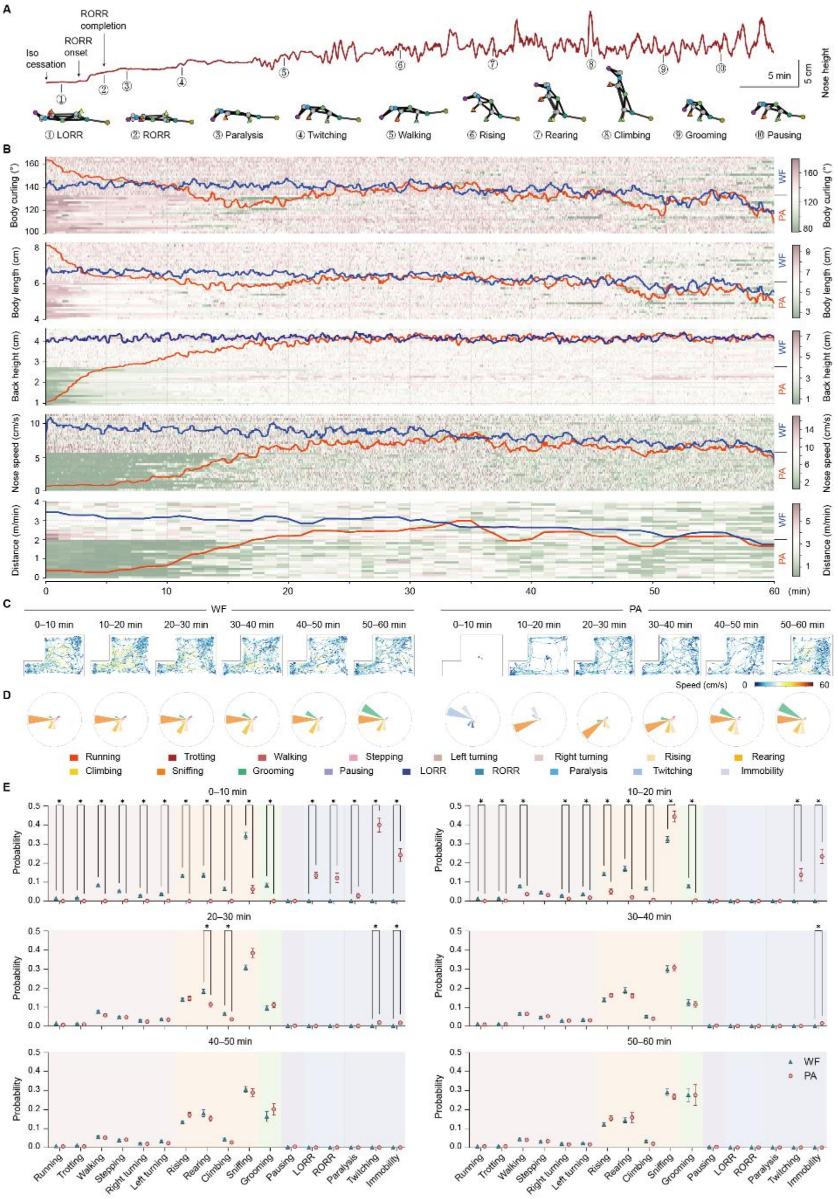


**Figure S2. Kinematic and behavioral dynamics during consciousness recovery , related to Figure 1**

(A) Representative nasal height dynamics and postural states following anesthesia cessation.

(B) Key kinematic parameter trajectories during WF and PA periods (n = 9 male and 9 female mice).

(C) Representative spontaneous movement patterns across 10-minute epochs during WF (left) and PA (right) periods.

(D) Polar representations of movement repertoire across 10-minute epochs during WF (left) and PA (right).

(E) Comparative movement probabilities between states.

Data are represented as mean (B and D) and mean ± s.e.m. (E). Statistical analyses are performed using multiple Mann-Whitney tests with Bonferroni–Dunn correction for multiple comparisons. ***$P < 0.001$, **$P < 0.01$, *$P < 0.05$. Statistical details are presented in Table S2.

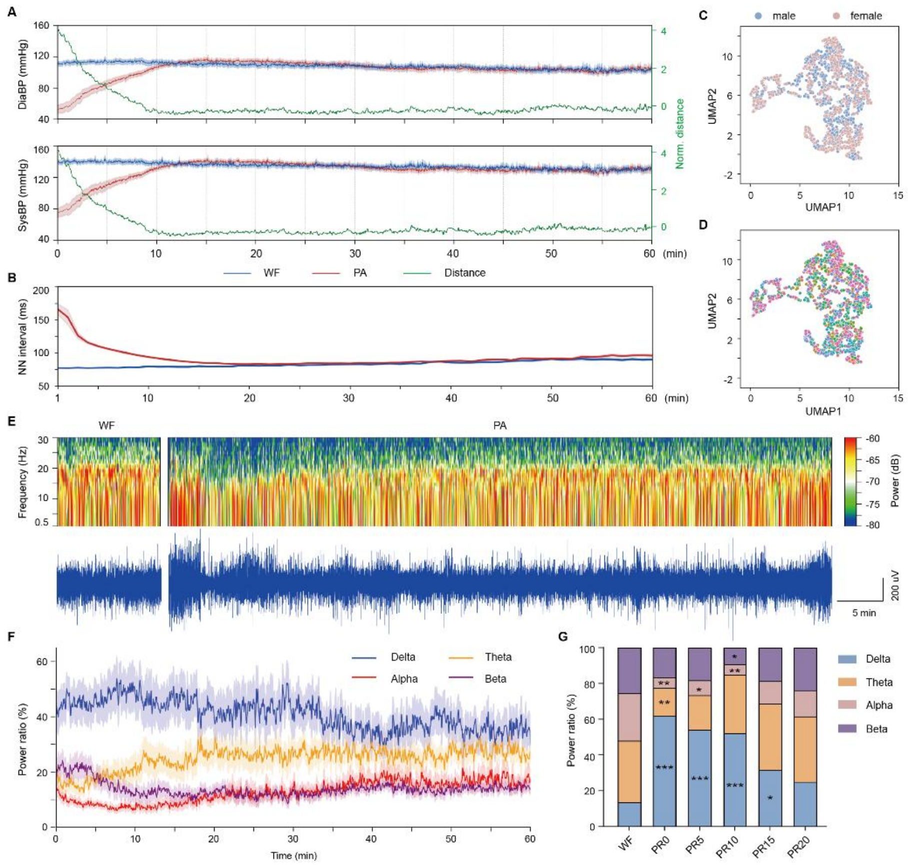


**Figure S3. Cardiovascular and EEG recovery signatures, related to Figure 1**

(A) Diastolic blood pressure (DiaBP, top) and systolic blood pressure (DiaBP, bottom) during WF and PA 60-minute period (left Y-axis), and DTW distance between WF and PA sequences for Dia BP and Sys BP (right Y-axis).

(B) NN interval progression during recovery.

(C and D) Sexes (C) and individuals (D) consistency in multimodal recovery trajectories.

(E) Representative EEG power spectrograms (top) and traces (bottom). (F) Spectral power band dynamics.

(G) Quantitative EEG band comparisons (n = 4 male and 4 female mice).

Data are represented as mean (G) and mean ± s.e.m. (A, B, and F). Statistical analyses are performed using Mann-Whitney tests with Bonferroni–Dunn correction for multiple comparisons. ***$P < 0.001$, **$P < 0.01$, *$P < 0.05$. Statistical details are presented in Table S2

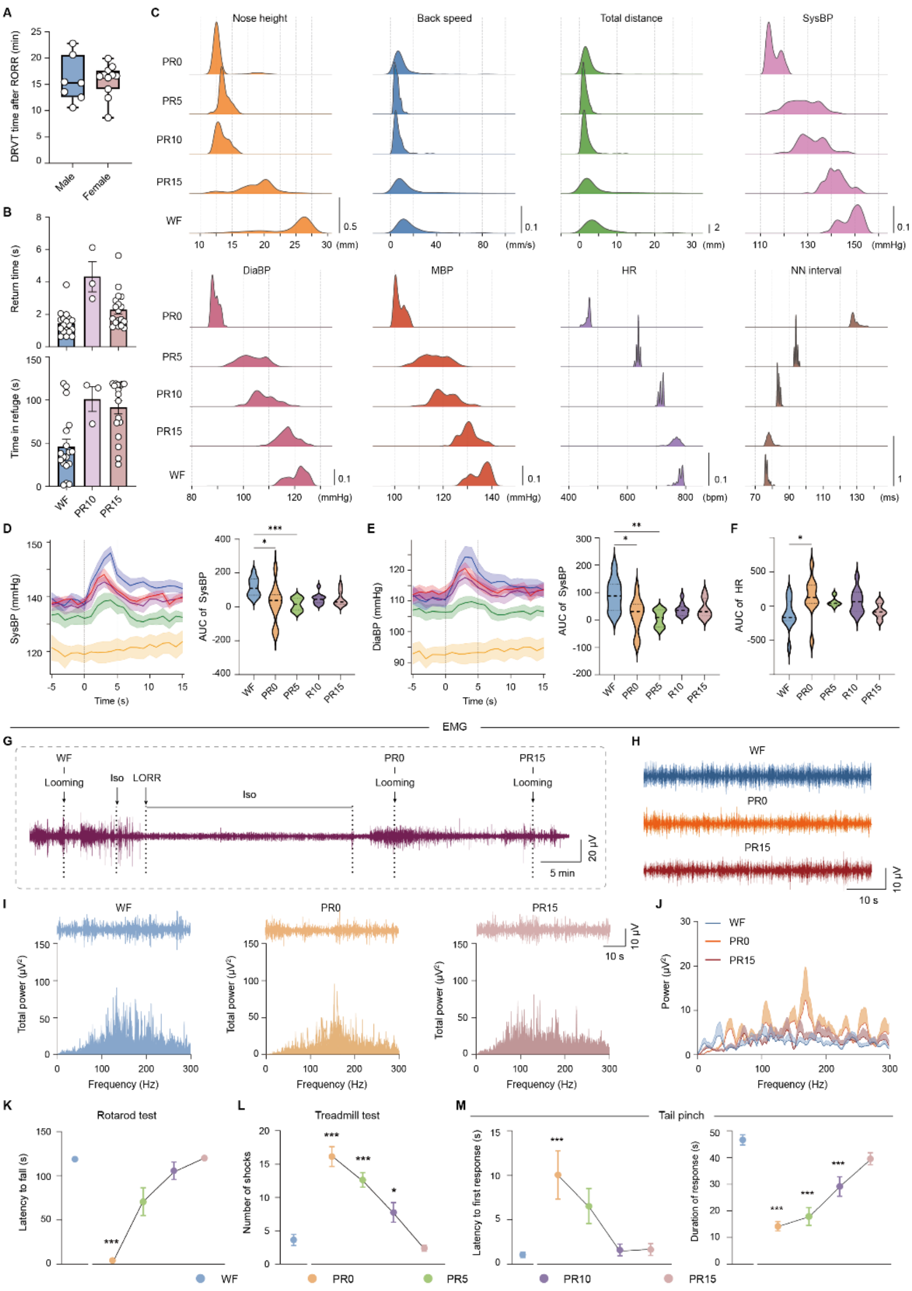


**Figure S4. Multimodal profiling of defensive behavior recovery, related to Figure 2**

(A) DRVT time after RORR in male (n =7) and female (n =11) mice.

(B) Return time (top) and time in refuge (bottom) comparisons.

(C), Ridgeline distributions of kinematic and cardiovascular parameters.

(D and E) SysBP (D) and DiaBP (E) trajectories (left) and AUC (right). (F) Heart rate response magnitudes.

(G) Representative EMG trace during WF-anesthesia-PA transition.

(H) Averaged EMG responses post-looming (n = 4 male and 4 female mice).

(I) Representative EMG traces (top) and spectral profiles (bottom) within 1 minute.

(J) EMG spectral power analysis within 1 minute. The trajectories were smoothed by 10 neighbors on each size.

(K and L) Rotarod (K) and treadmill (L) performance across timepoints (n = 4 male and 4 female mice).

(M) Nociceptive response latency (left) and durations (right) across consciousness timepoints (n = 5 male and 5 female mice).

Data are represented as mean ± s.e.m. (A; B; D and E, left; K–M), median with quartiles (D and E, right; F), and probability density (C). Statistical analyses are performed using unpaired t-test (A), Kruskal-Wallis test with Dunn's multiple comparisons test (B, D to F, and K), and one-way ANOVA with Dunnett's post hoc test (L and M). ***$P < 0.001$, **$P < 0.01$, *$P < 0.05$. Statistical details are presented in Table S3.

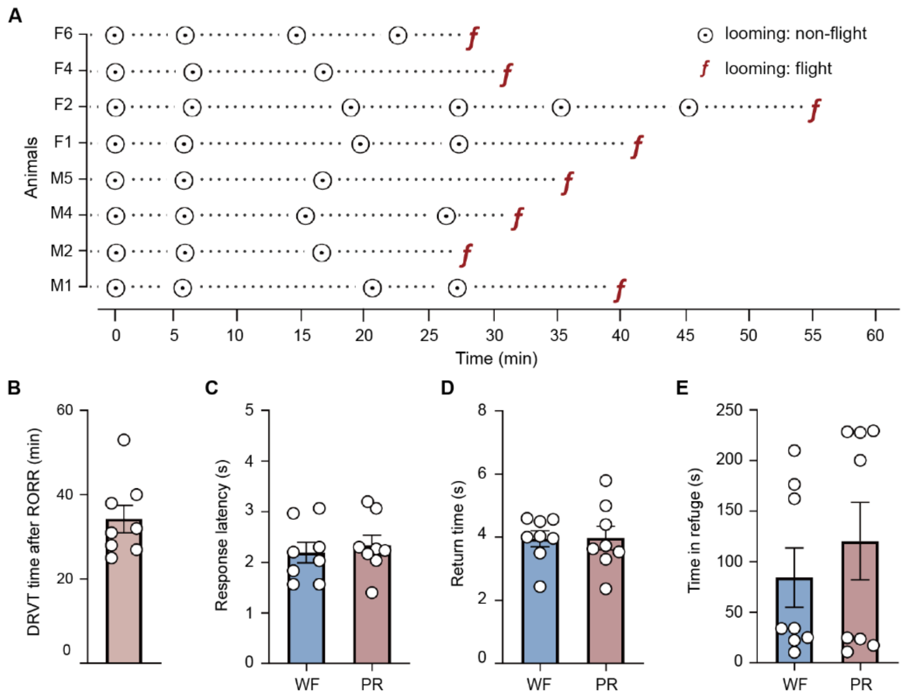


**Figure S5. Anesthetic-independent consciousness level recovery, related to Figure 2**

(A) Individual trails of responses to looming stimuli during propofol recovery after propofol discontinuation (n = 4 male and 4 female mice).

(B) Mean DRVT recovery time after RORR.

(C to E) Comparison of responsive latency (C), return time (D), and time in refuge (E). Data represent mean ± s.e.m. (box plots).

Data are represented as mean ± SEM. Statistical analyses are performed using unpaired t-test. $***P < 0.001$, $**P < 0.01$, $*P < 0.05$. Statistical details are presented in Table S3.

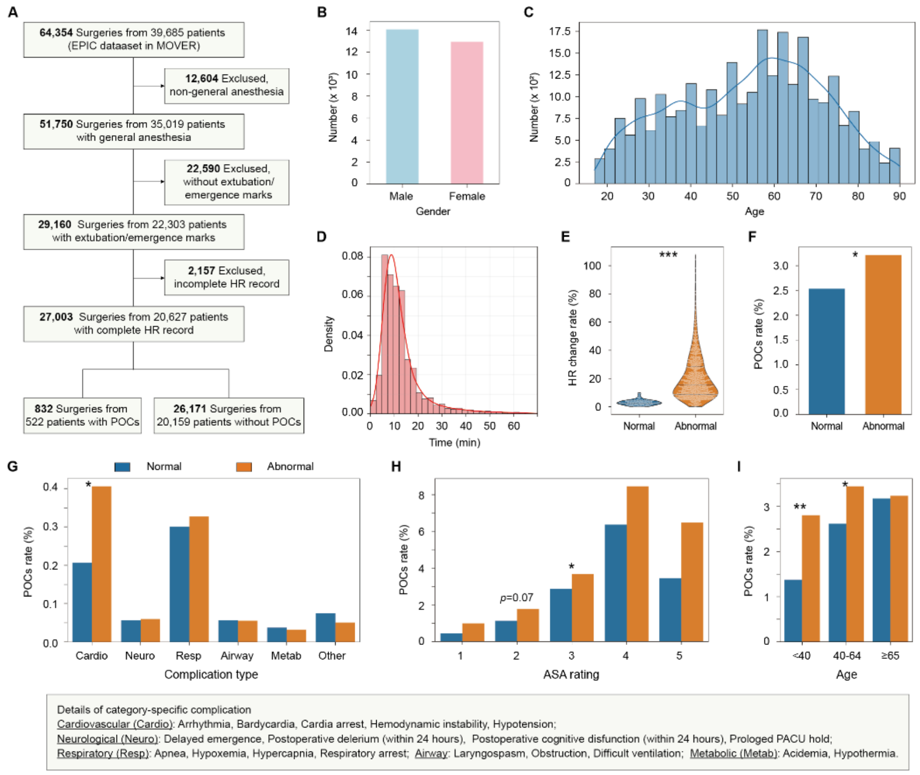


**Figure S6. Association between heart rate recovery patterns and postoperative complications in the MOVER dataset, related to Figure 2**

(A) Study flowchart for patient selection from the EPIC dataset in MOVER. Cases were excluded for: non-general anesthesia (n = 12,604), missing extubation/emergence documentations (n = 22,590), and incomplete heart rate (HR) records (n = 2,157). Final cohort included 27,003 analyzable cases from 20,627 patients (832 cases with postoperative complications (POCs); 26,171 cases without POCs).

(B and C) Gender (B) and age (C) distributions of the selected 27,003 cases.

(D) Density distribution of time difference between postoperative HR records and the closest key anesthetic events (extubation or emergence).

(E) HR change distribution in POCs cases. Abnormal recovery was defined as >10% deviation of postoperative HR from the preoperative baseline.

(F) Overall POCs rates stratified by HR recovery status.

(G) Category-specific complication characteristics. Cardiovascular complications showed strongest association with abnormal recovery. The details of complication types were described in the below rectangular box.

(H and I) POCs rates stratified by ASA physical status (H) and age groups (I).

Data are represented as percentages (F–I) and median with quartiles (E). Statistical analyses are performed using Fisher's exact test. ***$P < 0.001$, **$P < 0.01$, *$P < 0.05$. Statistical details are presented in Table S3.

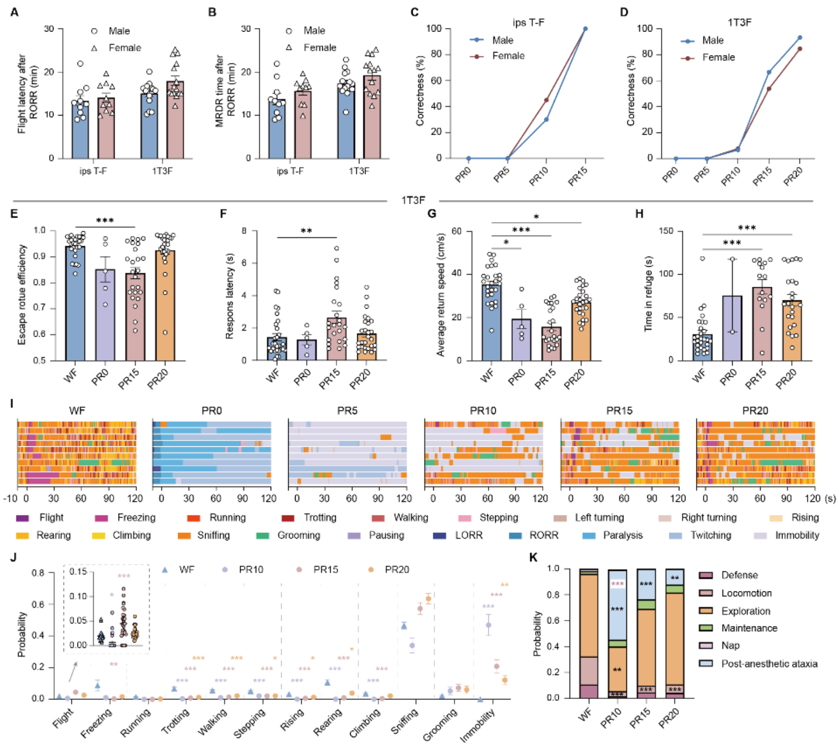


**Figure S7. Defensive behavior metrics of consciousness content recovery, related to Figure 3**

(A and B) Latency of flight initiation (A) and MRDR recovery (B) after RORR in male and female mice (n = 20 mice (10 per sex) in ips T-F, n = 29 mice (15 male and 14 female) in 1T-3F).

(C and D) Escape accuracy in ips T-F (C) and 1T-3F (D) conditions in male and female mice.

(E–H) Temporal evolution of escape route efficiency (E), response latency (F), return speed (G), and time in refuge (H) in 1T-3F configuration. (I) Representative ethogram across timepoints.

(J and K) Movement probability (J) and behavioral cluster distribution (K) comparisons. Data are represented as mean (C, D, and K) and mean ± s.e.m. (A, B, E–H, and J). Statistical analyses are performed using two-way ANOVA test with Bonferroni's post hoc test (A and B), Kruskal-Wallis test with Dunn's multiple comparisons test (E to H), and Mann-Whitney tests with Bonferroni–Dunn correction for multiple comparisons (J and K). ****P* < 0.001, ***P* < 0.01, **P* < 0.05. Statistical details are presented in Table S4.

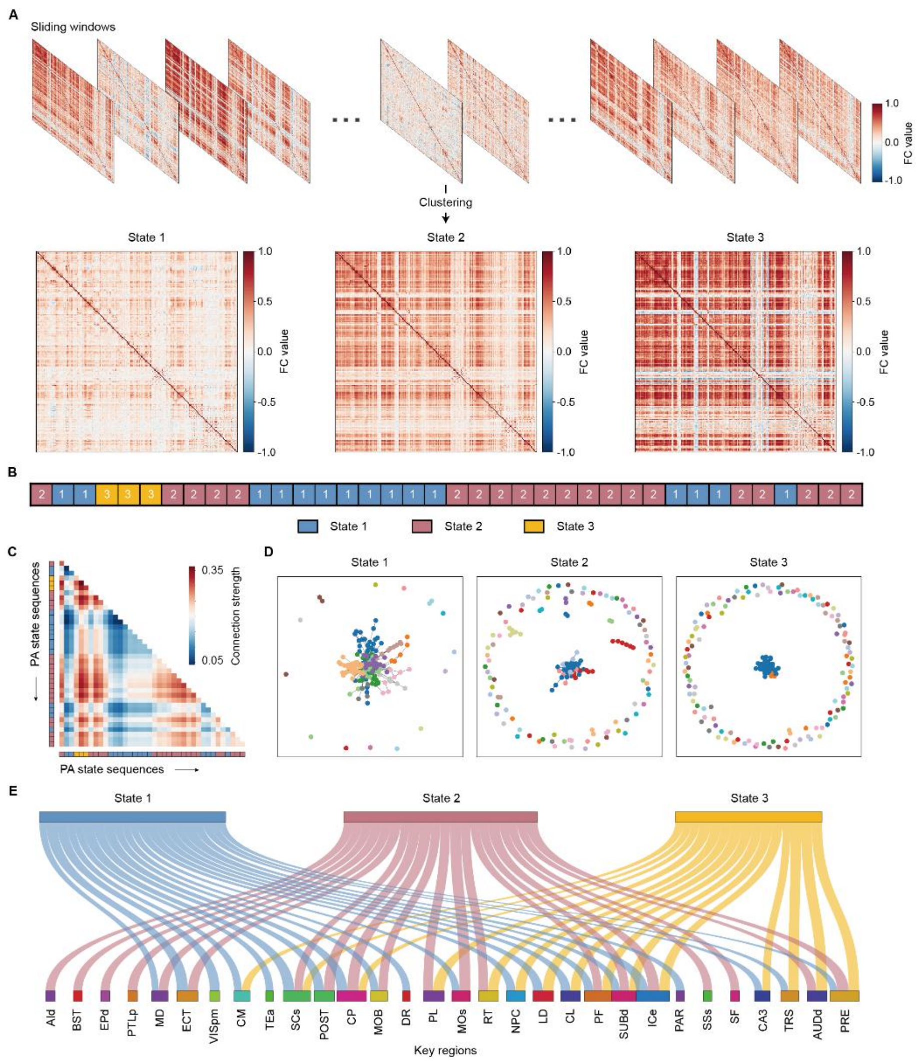


**Figure S8. Dynamic functional connectivity analysis framework, related to Figure 4**

(A) Sliding window approach schematic (top) and state-specific average FC matrices (bottom).

(B) Schematic of state sequences across time windows.

(C) Heatmap of FC strength across PA state sequences.

(D) Topology of each brain state.

(E) Region (bottom)-state (top) association patterns.

Anatomic abbreviations are shown in Table S5.

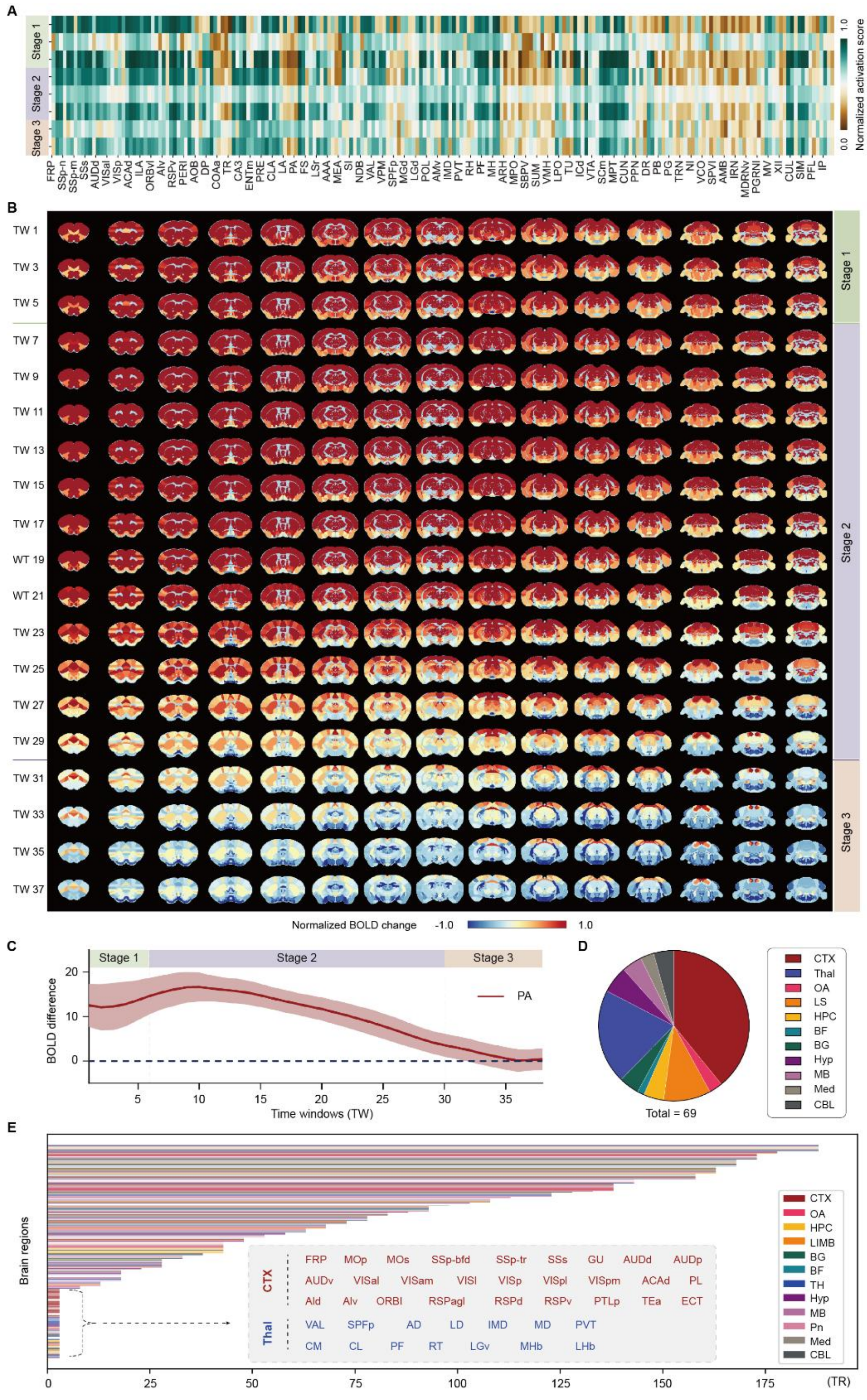


**Figure S9. Spatiotemporal BOLD dynamics and recovery initiation across brain regions, related to Figure 4**

(A) Normalized activation score of 214 brain regions (Allen Mouse Brain Atlas) across stages.

(B) Spatiotemporal evolution of Blood-oxygen-level-dependent (BOLD) changes. The images were presented at an interval of every other time window.

(C) BOLD signal differences relative to WF.

(D) Brain regions that initiate recovery earliest.

(E) Temporal sequence of recovery initiation across brain regions. The bracket indicates the brain regions with the earliest recovery onset. The rectangle specifically outlines the cortical and thalamic regions within this earliest-recovering group.

Data are represented as mean ± s.e.m. (C). Temporal sequences of recovery initiation across brain regions are shown in Table S5.

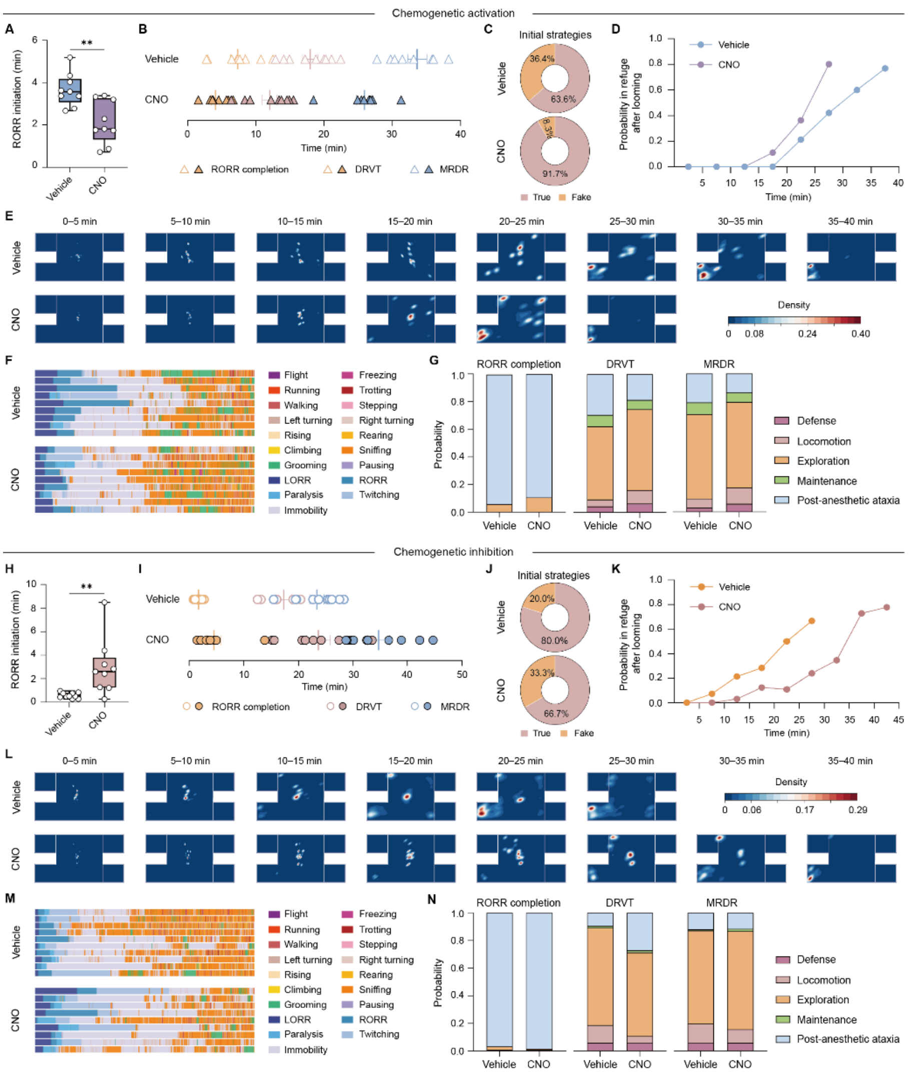


**Figure S10. Comprehensive recovery profiling following manipulation of the centromedian nucleus of thalamus, related to Figure 5**

(A and B) RORR initiation (A) and recovery landmark trajectories (B) following CM chemogenetic activation (n = 9 mice (5 male and 4 female) in per group).

(C) Escape strategy distributions following CM activation.

(D and E) Refuge probability (D) and spatial density (E) following CM activation.

(F and G) Behavioral ethogram (F) and behavior clusters distributions (G) following CM activation.

(H and I) RORR initiation (H) and recovery landmark trajectories (I) following CM chemogenetic inhibition (n = 10 mice (5 per sex) in vehicle and 9 mice (5 male and 4 female) in CNO).

(J) Escape strategy distributions following CM inhibition.

(K and L) Refuge probability (K) and spatial density distribution (L) following CM inhibition.

(M and N) Behavioral ethogram (M) and behavior clusters distributions (N) following CM inhibition.

Data are represented as mean (D, G, K, and N) and mean ± s.e.m. (A and H). Statistical analyses are performed using unpaired t-test (A and H) and Mann-Whitney tests with Bonferroni–Dunn correction for multiple comparisons (G and N). Statistical details are presented in Table S6.

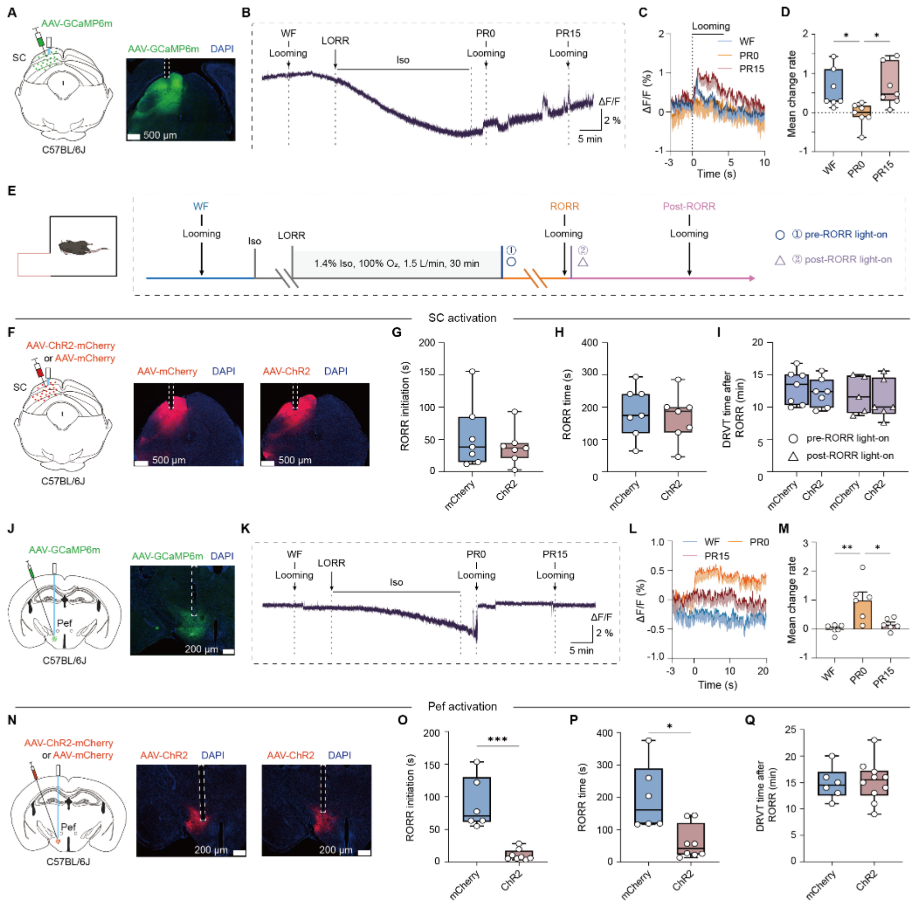


**Figure S11. Superior colliculus and perifornical area of the hypothalamus exhibit limited recovery promotion, related to Figure 5**

(A) Superior colliculus (SC) calcium imaging strategy and expression verification.

(B) Representative SC calcium dynamics.

(C and D) SC response profiles (C) and mean change rates (D) across timepoints (n = 4 male and 3 female mice).

(E) Optogenetic stimulation paradigms.

(F) SC optogenetic targeting.

(G–I) Time of RORR initiation (G), RORR completion (H), and DRVT after RORR (I) following SC activation (n = 7 mice (4 male and 3 female) per group).

(J) Schematic of viral injection and representative image of AAV-hSyn-GCaMP6m expression in the perifornical area of the hypothalamus (Pef) calcium imaging strategy and expression verification.

(K) Representative Pef calcium dynamics.

(L and M) Pef response profiles (L) and mean change rate (M) across timepoints (n = 3 male and 3 female mice).

(N) Pef optogenetic targeting.

(O–Q) Time of RORR initiation (O), RORR completion (P), and DRVT after RORR (Q) following Pef activation (n = 6 mice (3 per sex) in mCherry, n = 8 mice (4 per sex) in ChR2).

Scale bars, 500 μm (A and F), 200 μm (N). Data are represented as mean ± s.e.m. Statistical analyses are performed using one-way ANOVA (D and M) and two-way ANOVA (I) with Bonferroni's post hoc test, Mann-Whitney U test (O and P), and unpaired t-test (G, H, and Q). ***$P < 0.001$, **$P < 0.01$, *$P < 0.05$. Statistical details are presented in Table S6.
